# Next-generation intranasal Covid-19 vaccine: a polymersome-based protein subunit formulation that provides robust protection against multiple variants of concern and early reduction in viral load of the upper airway in the golden Syrian hamster model

**DOI:** 10.1101/2022.02.12.480188

**Authors:** Jian Hang Lam, Devendra Shivhare, Teck Wan Chia, Suet Li Chew, Gaurav Sinsinbar, Ting Yan Aw, Siamy Wong, Shrinivas Venkatraman, Francesca Wei Inng Lim, Pierre Vandepapeliere, Madhavan Nallani

## Abstract

Severe acute respiratory syndrome coronavirus 2 (SARS-CoV-2) is the etiological agent of coronavirus disease 2019 (Covid-19), an ongoing global public health emergency. Despite the availability of safe and efficacious vaccines, achieving herd immunity remains a challenge due in part to rapid viral evolution. Multiple variants of concern (VOCs) have emerged, the latest being the heavily mutated Omicron, which exhibits the highest resistance to neutralizing antibodies from past vaccination or infection. Currently approved vaccines generate robust systemic immunity, yet poor immunity at the respiratory tract. We have demonstrated that a polymersome-based protein subunit vaccine with wild type (WT) spike protein and CpG adjuvant induces robust systemic immunity (humoral and T cell responses) in mice. Both antigen and adjuvant are encapsulated in artificial cell membrane (ACM) polymersomes – synthetic, nanoscale vesicles that substantially enhance the immune response through efficient delivery to dendritic cells. In the present study, we have formulated a vaccine candidate with the spike protein from Beta variant and assessed its immunogenicity in golden Syrian hamsters. Two doses of ACM-Beta spike vaccine administered via intramuscular (IM) injection evoke modest serum neutralizing titers that are equally efficacious towards WT and Beta viruses. In contrast, the ACM-WT spike vaccine induces a predominantly WT-specific serum neutralizing response with pronounced reduction in potency towards the Beta variant. Remarkably, immunogenicity of the ACM-Beta spike vaccine is greatly enhanced through intranasal (IN) administration. Following IN challenge with the Beta variant, IM-immunized hamsters are fully protected from disease but not infection, displaying similar peak viral RNA loads in oral swabs as non-vaccinated controls. In contrast, hamsters IN vaccinated with ACM-Beta spike vaccine are protected from disease and infection, exhibiting a ∼100-fold drop in total and subgenomic RNA load as early as day 2 post challenge. We further demonstrate that nasal washes from IN-but not IM-immunized animals possess virus neutralizing activity that is broadly efficacious towards Delta and Omicron variants. Altogether, our results show IN administration of ACM-Beta spike vaccine to evoke systemic and mucosal antibodies that cross-neutralize multiple SARS-CoV-2 VOCs. Our work supports IN administration of ACM-Beta spike vaccine for a next-generation vaccination strategy that not only protects against disease but also an infection of the respiratory tract, thus potentially preventing asymptomatic transmission.

## Introduction

Severe acute respiratory syndrome coronavirus 2 (SARS-CoV-2), the etiological agent of coronavirus disease 2019 (Covid-19), has spread rapidly worldwide since its discovery in December 2019 and continues to drive a global public health emergency. At the time of writing (8^th^ February 2022), the World Health Organization (WHO) has reported more than 394 million infections and more than 5.7 million deaths globally (https://covid19.who.int/). Vaccination represents the best form of defence against SARS-CoV-2 and the world is pursuing vaccine research and development at an aggressive pace. Around 180 candidates of various modalities are in clinical development, of which approximately 80% are based on the viral spike (https://covid19.trackvaccines.org/agency/who/).

The spike glycoprotein forms homotrimers on the virion surface and is essential for viral entry into the host cell. Each monomer comprises of S1 and S2 domains, which serve distinct and critical roles during the cell entry process ^1^. Notably, the S1 subunit contains the receptor-binding domain (RBD) which binds the host cell angiotensin converting enzyme 2 (ACE2) receptor. This causes the spike protein to transit from a metastable prefusion state to a more stable post-fusion state, which is required for the fusion of virus and host cell membranes ^2^. Given the surface-exposed nature of the spike protein, it is the main target of neutralizing antibodies which predominantly recognize the RBD ^3, 4^, though recognition of S1 N-terminal domain (NTD) and the S2 subunit has also been reported ^5–8^. Importantly, the most potent neutralizing activity is associated with antibodies that directly block RBD interaction with ACE2 ^9^. Consequently, the spike protein has attracted intense attention as a target for vaccine development.

Multiple SARS-CoV-2 variants have emerged from rapid viral evolution, among which five variants of concern (VOCs) have been identified (https://www.who.int/en/activities/tracking-SARS-CoV-2-variants/): Alpha (B.1.1.7), first detected in the United Kingdom in September 2020; Beta (B.1.351) in South Africa, May 2020; Gamma (P.1) in Brazil, November 2020; Delta (B.1.617.2) in India, October 2020; and Omicron (B.1.1.529) in Southern Africa, November 2021. VOCs are characterized by one or more of the following: increased transmissibility ^10, 11^, increased virulence ^12^, or resistance to neutralizing antibodies raised against the wild type (WT) virus ^13–15^. These features typically arise from the mutation of key residues within the spike protein. For instance, residue E484 within the RBD constitutes a dominant neutralizing epitope and is a site of principal importance. Deep mutation scanning and escape mutation experiments have revealed that substitution with K, Q, P, A, D or G is associated with severe reduction in neutralizing potency of polyclonal human immune serum or plasma ^16, 17^. Notably, E484K mutation is found in Beta and Gamma variants, and E484A in Omicron, which co-occurs with K417N/T and N501Y mutations. Mutation at position 417 confers modest resistance to neutralization ^4, 16^ whereas N501Y enhances ACE2 binding which may account for increased transmissibility ^18^. The Omicron variant, in particular, has acquired more than 30 mutations in the spike protein and these are believed to enable extensive antibody escape from previously infected or vaccinated individuals ^15, 19^. Despite the reduction in neutralizing potency, mRNA (mRNA-1273; BNT162b2) and adenovirus vector (ChAdOx-1 S; Ad26.COV2.S) vaccines based on the WT spike remain effective against VOCs for the prevention of severe disease and death ^20, 21^, possibly due to preservation of T cell reactivity towards conserved epitopes among variant spike proteins ^22–24^. Nevertheless, to address rapidly waning antibodies six months after mRNA vaccination ^25, 26^, and the extensive loss of neutralizing activity towards the Omicron variant, boosters are required to restore protection ^15, 27, 28^.

It is desirable for a vaccine to induce broad neutralizing responses towards all VOCs. Interestingly, individuals who are infected by the Beta variant subsequently develop a vigorous antibody response that cross-neutralizes, to varying extents, Alpha, Gamma and Delta variants ^14, 29–31^. Notably, major developers Pfizer/BioNTech, Moderna and AstraZeneca have initiated clinical trials to assess the safety and efficacy of vaccines based on the Beta variant spike ^32^. Against the highly mutated Omicron variant, however, neutralizing potency of Beta convalescent sera is greatly diminished ^29, 33^.

We have previously reported the preclinical development of a Covid-19 subunit spike protein vaccine based on our proprietary Artificial Cell Membrane (ACM) polymersome technology ^34^. These synthetic nanovesicles measure 100-200 nm in diameter and are made up of an amphiphilic block copolymer comprising of poly(butadiene)-b-poly(ethylene glycol) (PBD-*b*-PEO) and a cationic lipid, 1,2-dioleoyl-3-trimethylammonium-propane (DOTAP). In aqueous medium, the amphiphilic molecules self-assemble into a bilayer membrane vesicle that encapsulates soluble materials within its cavity. Functionally, ACM polymersomes are strongly taken up by antigen-presenting cells (APCs), including dendritic cells (DCs), thus enabling efficient delivery of encapsulated antigens and adjuvants for enhanced immunogenicity. Our Covid-19 vaccine is formulated with recombinant SARS-Cov-2 spike protein (hereby referred to as “S1S2”) and CpG adjuvant separately encapsulated in ACM polymersomes for co-administration. Such approach confers modularity and flexibility in the choice of spike protein to include in the formulation, which is highly relevant given the rapid emergence of new VOCs. Moreover, we and others ^34, 35^ have shown that adjuvant effect of CpG is not impaired despite the lack of physical linkage to the spike protein. Mice co-administered with ACM-S1S2 + ACM-CpG developed strong, durable serum neutralizing titers and functional memory T cells, whereas mice immunized with the non-encapsulated formulation generated weaker responses that rapidly waned ^34^. In the present study, we investigate the immunogenicity of ACM Covid-19 vaccine based on WT or Beta spike in golden Syrian hamsters, followed by its ability to protect against Beta variant challenge. Hamsters are naturally susceptible to SARS-CoV-2 and can support viral replication as well as transmission, making it an important small animal model for studying pathogenesis as well as efficacy of prophylactics and therapeutics ^36, 37^. We show that intramuscular (IM) administration with an ACM vaccine based on WT spike generates antibodies that predominantly neutralize WT virus, whereas another ACM vaccine based on Beta spike induces antibodies that cross-neutralize WT and Beta viruses. Both vaccines strongly protect against disease after intranasal (IN) challenge with the Beta variant but do not efficiently suppress infection of the upper respiratory tract. Remarkably, immunogenicity of the ACM-Beta spike vaccine is strongly enhanced by IN administration. Moreover, IN vaccination potently reduces viral RNA load by ∼100-folds as early as Day 2 post challenge in the nasal swabs. Accordingly, we detect virus neutralizing activity in the nasal washes of IN-but not IM-vaccinated hamsters, indicating a protective mucosal immune response that correlates with suppression of viral replication in the upper respiratory tract. Antibodies induced by two IN doses of ACM-Beta spike vaccine are broadly neutralizing and retain appreciable activity towards Omicron, suggesting that our mucosal vaccination strategy may protect against infection by this current variant and, potentially, future VOCs.

## Materials and methods

### Materials

Human CpG 7909 (T*C*G*T*C*G*T*T*T*T*G*T*C*G*T*T*T*T*G*T*C*G* T*T; where * denotes phosphorothioate bond) was synthesized by BioSpring. 1,2-dioleoyl-3-trimethylammonium-propane (DOTAP) was from Avanti Polar Lipids. Triton X-100 was from MP Biomedicals. All other chemicals were purchased from Sigma-Aldrich unless stated otherwise.

### Baculovirus production

The Beta variant spike protein ectodomain gene (amino acids 1-1201) containing the native signal peptide, 3Q mutations to the furin cleavage site and 2P mutations, was codon-optimized for insect cell protein expression, using GenScript proprietary algorithm and was directly synthesized into pFAST-BAC1 transfer plasmid. This transfer plasmid (500 ng) was transformed using heat shock (42 °C, 1 min) into competent DH10 BAC cells (Thermo Fisher Scientific). Cells were cultured on agar plates containing selection antibiotics and Blu-gal. Colonies that were positive for recombination were selected and Bacmid DNA was extracted using traditional Midiprep technology. In brief, a 50 ml culture was inoculated from the plate and grown for 16 hours at 37 °C at 200 rpm. The cells were pelleted, and the supernatant was removed. Pellet was resuspended into the resuspension buffer (25 mM Tris-HCl pH 8, 10mM EDTA, 100 µg/ml RNase) and allowed to incubate at room temperature for 5 minutes, followed by incubation with lysis buffer (0.2 M NaOH and 1% SDS). Finally, precipitation buffer (3 M potassium acetate, pH 5.5) was added, and the solution was left to incubate for a further 10 minutes at 4 °C, followed by centrifugation at 10,000 g, 4 °C, 15 minutes. The DNA was precipitated from the supernatant with 100% isopropanol and washed with 70% ethanol. The pellet was dried and finally resuspended in TE buffer pH 8.0. This DNA was used for transfection using the methods described in the Bac-to-Bac (Thermo Fisher Scientific) with Trans IT-Virus GEN (Mirus). In brief, 2.4 x10^5^ Sf9 insect cells were plated into a well of a 24 well plate. After 1 hour, the media was aspirated. A premix of transfection reagent and 200 ng of DNA was allowed to stand for 30 minutes at room temperature before being added to the plated cells. The plate was left to incubate for 6 hours followed by addition of the ESF-AF medium (Expression systems). Sf9 cells were left for seven days at 27 °C without agitation. The baculovirus-containing supernatant was collected and used to infect further cultures of Sf9 cells to amplify various recombinant virus stocks (P1-P3). Finally, P3 virus stock was titrated as described ^38^ and used for protein expression.

### Protein expression

Sf9 cells (Thermo Fisher Scientific) were routinely grown in ESF-AF medium and maintained at a cell density between 1-4 x 10^6^ cells/ml. For protein production, cells were adjusted to ∼2 x 10^6^ per ml in 600 ml culture volume, in a plastic 2 L non-baffled flask. The culture was incubated to a density of 2.5-3 x 10^6^ cells/ml and then infected with recombinant baculovirus (P3) at a multiplicity of infection (MOI) of 0.1. Flasks were left for 68-72 hours at 27 °C with shaking at 115 rpm. To harvest spike protein secreted into culture supernatant, cells were removed by centrifuging in a swing bucket rotor at 3,000 g, 10 mins, 4°C.

### Concentration and buffer exchange

A 50 kDa cut-off Tangential flow filtration column (Repligen) was washed and equilibrated as per manufacturer’s instructions. The harvested cell culture supernatant was then loaded onto the column and the retentate recirculated at a flow rate of 50 ml per minute. A 6-8 PSI transmembrane pressure was applied, and the sample was concentrated 10-fold and dia-filtered 5 times with the buffer 1 (20 mM phosphate, 100 mM NaCl, 5 % glycerol, pH 5). The final sample was centrifuged at 16,000 g for 15 minutes at 4 °C. The supernatant was filtered through a 0.22 µm PES filter before subsequent purification steps.

### Protein purification

1 x 5 ml GE Hitrap SP FF column and 1 x 5 ml GE Hitrap Q HP column were used. Each column was pre-equilibrated using a GE AKTA FPLC Explorer with their respective binding buffer i.e.: 20 mM phosphate, 100 mM NaCl, 5 % glycerol, pH 5 (Buffer 1) for SP, and 20 mM phosphate, 100 mM NaCl, 5 % glycerol, pH 8 (Buffer 2) for Q column, with a flowrate of 2 ml/min, for 10 column volumes (CV). The spike protein sample was placed on ice and loaded onto the SP column at 2 ml per minute until the entire sample was loaded. The unbound sample was collected for subsequent analysis. The column was washed with 10 CV of Buffer 1. After this, the bound protein on SP was eluted with buffer 2 by switching the pH. The protein from this SP elution step was collected for further purification using Q column. After 10 column volumes, the SP column was further eluted with buffer 3 (20 mM phosphate, 1M NaCl, 5 % glycerol, pH 8), sample was collected for later analysis. Finally, the SP column was washed with 5 CV of 0.1 N NaOH, sample was collected for later analysis and stored. In the next step, buffer 2 eluate from SP column was loaded on to the pre-equilibrated Q column until entire sample was loaded. Following loading, Q column was washed with buffer 2 for another 10 column volumes. After the Q wash, a gradient between buffer 2 and buffer 3 was set from 0% to 32% for 75 minutes at 2 ml per minute. 5 ml fractions were collected across the linear gradients. Recombinant Beta variant spike (S1S2) protein eluted at a conductivity of 33-37 mS/cm (between 26-30% B).

### Preparation of ACM-antigen polymersomes

ACM polymersomes encapsulating S1S2 protein was prepared by the solvent dispersion method, followed by extrusion. A 380 mg/ml stock solution of DOTAP and PEG_13_-*b*-PBD_22_ polymer were prepared by dissolving solid DOTAP and polymer in tetrahydrofuran (THF) to prepare solution A as described earlier ^34^.A 5 ml solution of 600 µg/ml S1S2 protein was placed in a 50 ml Falcon tube (Solution B). Solution A was added slowly to 5 ml of Solution B while constantly mixing (600-700 rpm) at room temperature. A turbid solution was obtained. The resulting solution was extruded 21 times through a 200 nm membrane filter (Avanti Polar Lipids) using a 1 ml mini-extruder (Avanti Polar Lipids) to get monodispersed ACM-antigen vesicles. Non-encapsulated antigens were removed by 2 days of overnight dialysis with 3 buffer exchanges. Encapsulation of antigen were quantified by densiometric analysis using a known S1S2 protein standards in Fiji ImageJ software (v. 1.52a).

### Preparation of ACM-CpG polymersomes

ACM-CpG polymersomes were prepared by the solvent dispersion method described above, followed by extrusion. 1.2 ml of the 700 mg/ml stock solution containing DOTAP and PEG_13_-*b*-PBD_22_ polymer was added dropwise to 10 ml CpG solution. A turbid solution was obtained. The resulting solution was extruded 21 times through a 200-nm membrane filter using a 1 ml mini-extruder to get monodispersed ACM-CpG polymersomes. Unencapsulated CpG was removed by overnight dialysis using 300 kDa molecular weight cut-off (MWCO) as described earlier ^34^.

### Particle size measurement by dynamic light scattering (DLS)

DLS was performed on the Zetasizer Nano ZS system (Malvern Panalytical) as previously described ^34^.

### SDS-PAGE

The protein quantity of encapsulated S1S2 protein was determined by the densitometry using SDS-PAGE gel as reported earlier ^34^. Briefly, protein encapsulated polymersomes were released with Triton X-100. 5 µl of 4x gel loading buffer was added into 15 µl of the reconstituted sample. The sample was vortexed and heated at 95 °C for 10 minutes. The heated sample was loaded into the 8% Bis-Tris gel and the gel was stained using SYPRO™ Ruby protein gel stain. The gel image was captured by the ImageQuant™ LAS 500 imager and analyzed using ImageJ software.

### Quantification of CpG

The CpG quantity of the encapsulated CpG was measured by the reversed-phase high-performance liquid chromatography (RP-HPLC). 15 µl of 10% Triton X-100 solution was added into 30 µl of sample to lyse the vesicles, followed by adding 105 µl of 100mM triethylammonium acetate (TEAA) buffer, pH 7.0. The mixture was vortexed, centrifuged and transferred into a 250 µl glass insert in 2mL HPLC vial and capped. The analytes were injected into Agilent AdvancedBio Oligonucleotides column with Agilent 1260 Series Capillary LC system. The UV intensity at wavelength 260 nm of the analytes was measured and the area under the peak was recorded for analysis.

### Assessment of free spike protein function by ACE2-binding assay

96-well high-binding EIA/RIA plates (Corning) were precoated overnight with spike protein at 200 ng per well. Wells were washed four times with Wash Buffer (TBS supplemented with 0.05% v/v Tween-20) and blocked with 2% w/v BSA in Wash Buffer for 1.5 hour at 37 °C. ACE2 protein conjugated to human Fc tag was three-fold serially diluted (dynamic range: 12,000 – 0.61 ng/ml) with 0.5% w/v BSA in Wash Buffer and 100 µl of each dilution was applied to the respective well of the ELISA plate. The plate was incubated 1 hour at 37 °C and then washed four times. To detect bound ACE2 protein, HRP-conjugated goat anti-human IgG (Fc) (Bio Rad) was applied at 1:10,000 and incubated for 1 hour at 37 °C. The plate was washed four times before TMB (Sigma Aldrich) was added. After 10 minutes of color development, the reaction was terminated with equal volume of ELISA Stop Solution (Thermo Fisher Scientific). Absorbance at 450 nm was read using the BioTek plate reader. Background absorbance was subtracted and the EC_50_ value of the titration curve was determined using GraphPad Prism version 8.4.3 with five-parameter non-linear regression.

### Assessment of encapsulated spike protein function by ACE2-binding assay

Encapsulated spike protein was released by lysing ACM vesicles with 1% Triton-X 100 for 10 minutes at room temperature. Detergent was removed by frequent agitation with 200 mg/ml of Bio-Beads SM-2 Resin (Bio Rad) for 1 hour at room temperature. 96-well high-binding EIA/RIA plates (Corning) were precoated overnight with ACE2 protein at 500 ng per well. Wells were washed four times with Wash Buffer (TBS supplemented with 0.05% v/v Tween-20) and blocked with 2% w/v BSA in Wash Buffer for 2 hours at room temperature. Spike protein was three-fold serially diluted (dynamic range: 12,000 – 0.61 ng/ml) with 0.5% w/v BSA in Wash Buffer and 100 µl of each dilution was applied to the respective well of the ELISA plate. The plate was incubated 2 hours at room temperature and then washed four times. To detect bound spike protein, mouse anti-S1 mAb (clone 1035206; R&D Systems) was diluted 1:100 and applied to each well. The plate was incubated 2 hours at room temperature and then washed four times. Finally, HRP-conjugated goat anti-mouse IgG (H/L) (Bio Rad) was applied at 1:3,000 and incubated for 1 hour at room temperature. The plate was washed four times before TMB (Sigma Aldrich) was added. After 10 minutes of color development, the reaction was terminated with equal volume of ELISA Stop Solution (Thermo Fisher Scientific). Absorbance at 450 nm was read using the BioTek plate reader. Background absorbance was subtracted and the EC_50_ value of the titration curve was determined using GraphPad Prism version 8.4.3 with five-parameter non-linear regression.

### Hamsters (vaccination)

Animal procedures, sample collection and analyses were performed by Bioqual, Inc. (USA) under a paid service agreement. Golden Syrian hamsters comprising equal numbers of males and females were purchased from an approved vendor. Each group (n = 8) was administered a specific formulation via IM or IN route (see table below). Each dose was standardized at 20 µg spike protein and 100 µg CpG for a total of two doses, 21 days apart. Blood and nasal wash were collected 20 days after first dose and 13 days after second dose to assess antibody levels.

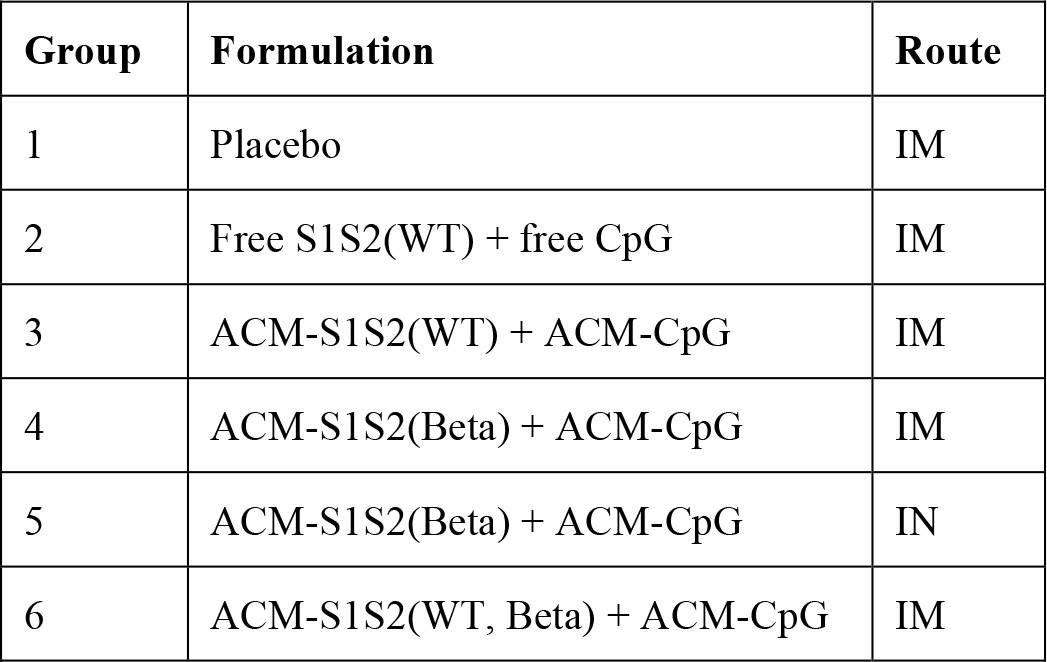

| Group | Formulation | Route |
| --- | --- | --- |
| 1 | Placebo | IM |
| 2 | Free S1S2(WT) + free CpG | IM |
| 3 | ACM-S1S2(WT) + ACM-CpG | IM |
| 4 | ACM-S1S2(Beta) + ACM-CpG | IM |
| 5 | ACM-S1S2(Beta) + ACM-CpG | IN |
| 6 | ACM-S1S2(WT, Beta) + ACM-CpG | IM |

### Serum processing

Blood was collected from the retro-orbital vein under sedation, in SST tubes (BD) and allowed to clot for 30 minutes to 1 hour at room temperature. Blood collection volumes were limited to 0.78 µl/g every 24 hours; 5.85 µl/g every 7 days; and 7.8µl/g every 14 days or more. The samples were centrifuged at 1,000-1,300 g for 5-10 minutes with brakes off. Serum was collected using a P200 pipettor, in a 0.5 ml cryovial and stored at −20 °C until analysis.

### Nasal wash

Hamsters were anesthetized using isoflurane, placed in lateral recumbency, and using a soft tipped catheter, 400 ul of PBS was flushed into the nasal cavity. A collection device was placed under the opposite nostril to collect the fluid (∼200-250 µl yield). The nasal wash was collected in a 2 ml cryovial and snap frozen.

### Hamsters (SARS-CoV-2 challenge)

14 days after final vaccination, hamsters were challenged with SARS-CoV-2 RSA (Beta variant; LOT 030621-750) at a dose of 3.67 x10^2^ PFU in 100 µl via IN inoculation. Prior to the procedure, animals were anesthetized by injecting 80 mg/kg ketamine and 5 mg/kg xylazine via IM route. Using a calibrated P200 pipettor, 50 μl of the viral inoculum were administered dropwise into each nostril. Anesthetized animals were held upright such that the nostrils of the hamsters were pointing towards the ceiling. The tip of the pipette was placed into the first nostril and virus inoculum slowly administered into the nasal passage, and then removed. This was repeated for the second nostril. The animal’s head was tilted back for about 20 seconds and then returned to its housing unit. Animals were injected with an antisedan for the reversal of anesthesia, at 1 mg/kg via IM route, 20 minutes post-challenge and monitored until complete recovery. Over the next 14 days, hamsters were monitored for weight changes, physical signs of distress and oral swabbed at designated time points for viral RNA qPCR.

### Oral Swabs

Hamsters were restrained properly for sample collection. A sterile swab was removed from the packaging, and the oral cavity was swabbed once. The swab was placed into a cryovial with 1ml PBS, the shaft cut off to fit into the vial, and the vial placed immediately on dry ice for snap-freezing. The sample was then stored at −80 °C until analysis.

### Serum IgG ELISA

Indirect ELISA was performed to analyze sera for binding antibodies to SARS-CoV-2 spike protein. Nunc MaxiSorp 96-well plates were coated with 100 µl of commercial recombinant spike (WT or Beta variant; Sino Biological) diluted to 2 µg/ml in PBS, pH 7.4. Plates were incubated statically for 12 hours at 37 °C. Unbound antigen was removed by washing three times with PBS + 0.05% v/v Tween-20. Plates were blocked in PBS + 5% w/v skim milk for 1 hour at 37 °C. Test and positive control samples were diluted in assay diluent (1% w/v skim milk in wash buffer) to an initial dilution of 1:20 followed by four-fold serial dilution. Once blocking was completed, blocking buffer was discarded and each serum sample was plated in duplicate. Plates were incubated for 2 hours at 37 °C statically, followed by three washes to remove unbound sera. Secondary detection antibody (goat anti-species-HRP IgG; Abcam) was added at a dilution of 1:10,000 in 100 µl per well. Plates were incubated for 30 minutes at room temperature statically, and unbound antibodies were subsequently removed as described above. To develop, 100 µl of 1-Step Ultra TMB substrate was added to each well. The reaction was stopped after ∼ 10 minutes with 50 µl TMB stop solution (SERA CARE). The plates were read within 30 minutes at 450 nm with a Thermo Labsystems Multiskan spectrophotometer. Antibody titer was defined as the reciprocal of the highest dilution that gave a pre-defined cut-off value for OD450. Serum titration curves were analyzed by 5-parameter, non-linear regression using GraphPad Prism version 8.4.3 to determine antibody titers.

### SARS-CoV-2 surrogate virus neutralization test (cPass™)

The cPass™ kit (GenScript) was used according to manufacturer’s instructions. Briefly, each sample was diluted 1:10 using Sample Dilution Buffer and incubated with an equal volume of HRP-RBD (WT or variant) reagent for 30 minutes at 37 °C. The Omicron RBD sequence was based on the most prevalent mutations and reflect the dominant Omicron strain ^15^. The mixture of serum and HRP-RBD was then applied to eight-well strips pre-coated with ACE2 protein for 15 minutes at 37 °C. Unbound RBD was washed off and RBD-ACE2 binding was visualized by addition of TMB substrate for 15 minutes at room temperature. Reaction was terminated using Stop Solution and absorbance was measured at 450 nm. Inhibition of RBD-ACE2 binding was calculated using the formula: 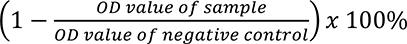. Where necessary, samples were pooled and concentrated using Vivaspin® 500 centrifugal concentrators (MWCO 50 kDa; Sartorius).

### Plaque reduction neutralization test (PRNT)

Vero 76 cells were cultured in 24-well plates at 175,000 cells/well in DMEM + 10% v/v FBS + Gentamicin and incubated at 37 °C, 5% CO_2_. Cells were used at 90-100% confluency. Serum samples were heat inactivated at 56 °C for 30 minutes. 30 PFU/well concentration of virus was prepared and kept on ice until use. Each serum sample was first diluted 1:10 with DMEM + 2% v/v FBS + gentamicin, followed by three-fold serial dilutions. Equal volume of 30 PFU/well virus inoculum was added to each serum dilution. Virus-only positive control and no-virus negative control were prepared in parallel. The mixes were incubated at 37 °C, 5% CO_2_ for 1 hour. Subsequently, Vero cell culture medium was removed from 24-well plate and 250 μl of titrated serum samples were added in duplicates. The 24-well plate was incubated at 37 °C, 5% CO_2_ for 1 hour for virus infection. During this time, 0.5% w/v methylcellulose medium was pre-warmed in a 37 °C water bath. Subsequently, 1 ml of methylcellulose medium was added to each well and the plate was incubated at 37 °C, 5 % CO_2_ for three days. The overlay medium was then removed, and Vero monolayers were washed once with 1 ml PBS. Cells were fixed with 400 μl ice-cold methanol per well at −20°C for 30 minutes. After fixation, methanol was discarded and the monolayers were incubated with 400 μl per well of staining solution (0.2% w/v crystal violet, 20% v/v methanol, 80% v/v dH_2_O) for 30 minutes at room temp. Wells were washed once with PBS or dH_2_O and allowed to dry for ∼15 minutes. The plaques in each well were recorded and the number of infectious units calculated.

### Viral RNA load determination by qPCR

The amount of RNA copies per ml of oral swab was determined using a validated qRT-PCR assay. Viral RNA was first isolated from samples using Qiagen DSP Virus/Pathogen Midi Kit and IVD Complex800 or IVD Cellfree500 protocols. The qRT-PCR assay utilized primers and a probe specifically designed to amplify and bind a conserved region of Envelop (E) gene of SARS-CoV-2 for the genomic RNA and Nucleocapsid (N) gene for sub-genomic RNA detection. The signal was compared to a known standard curve and calculated to give copies per ml. To generate a control for the amplification reaction, RNA was isolated from the applicable SARS-CoV-2 plasmid control using the same procedure. qPCR was set up using TaqMan Fast Virus 1-Step Real-time RT-PCR protocol with assay setup performed using Qiagen Qiagility automated PCR setup platform and analyzed in Applied Biosystems on QuantStudio 3.

### Statistics

Analyses were done using GraphPad Prism software (version 9.2.0). Where appropriate, one-or two-way repeated measures ANOVA (Greenhouse-Geisser correction) with Tukey’s multiple comparisons, or two-tailed paired T test was performed. Significant differences were indicated where present. *: *P* ≤ 0.05; **: *P* ≤ 0.01; ***: *P* ≤ 0.001; ****: *P* ≤ 0.0001; ns: not significant.

## Results

### Expression and purification of recombinant Beta and WT spike (S1S2) protein

Sf9 insect cells were transfected with recombinant baculovirus containing the gene encoding S1S2 ectodomain of SARS-CoV-2 from Beta variant (B.1.351). The gene was strategically modified to encode two consecutive proline substitutions in the S2 subunit in a turn between the central helix and heptad repeat 1 (HR1) to stabilize the pre-fusion conformation ^39^. This was considered desirable to generate relevant neutralizing antibodies ^40^. In addition, three glutamine substitutions at the furin cleavage site were introduced to prevent separation of S1 and S2 subunits by cellular proteases. The protein secreted into the culture supernatant was purified by sequential cation and anion exchange chromatography (Fig. 1a and b**, respectively)**. SDS-PAGE analysis of fractions collected over the purification process showed increasing purity of two closely migrating major bands at 150 kDa (Fig. 1c). Further analysis by western blot using a commercially available spike-specific polyclonal antibody (Fig. 1e) and our previous experience with purifying WT spike ^34^ enabled us to ascertain their identity as the protein-of-interest (S1S2 protein). Fractions with highest purity from the Q column were pooled. Subsequent SDS-PAGE analysis confirmed the presence of major bands at 150 kDa, as well as minor indistinct bands at 100 kDa and 75 kDa (Fig. 1d). Scanning densitometry consistently indicated a purity of >90%. Using a similar method, recombinant S1S2 protein from WT virus was also expressed and purified (Fig. 1e). Finally, a ACE2-binding assay was performed by immobilizing spike protein on the ELISA plate followed by titration of recombinant ACE2 protein (Fig. 1f). Both S1S2(WT) and S1S2(Beta) were functional and bound ACE2 with representative EC_50_ values of 25.0 and 18.4 ng/ml, respectively.

**Figure 1.**
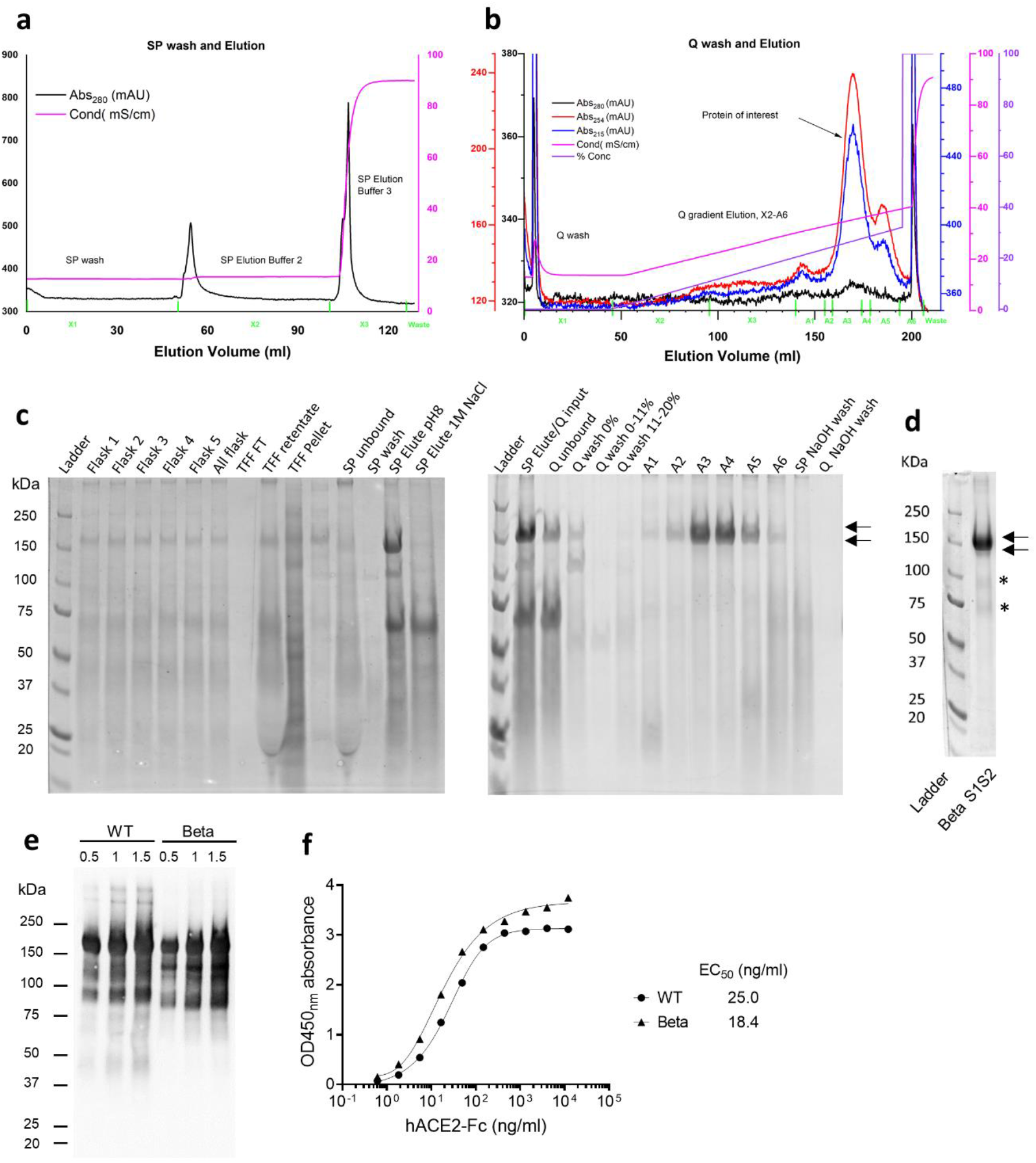
Purification of recombinant S1S2 protein (Beta variant). Sf9 insect cells were transfected with gene encoding S1S2 protein using recombinant baculovirus. Protein secreted into cell culture supernatant was purified by sequential passage through cation (SP) and anion (Q) exchange columns. **a.** Representative chromatogram of SP wash and eluate. **b.** Representative chromatogram of Q wash and eluate. (Blue) UV 280 mAU; (Red) UV 254 mAU; (Pink) UV 215 mAU; (Brown) conductivity mS/cm; (Green) percentage of Buffer 3. The initial loading of the sample is not shown. **c.** SDS-PAGE analysis of all fractions collected throughout the purification process. Protein bands were visualized by SYPRO Ruby stain. Arrows indicate protein-of-interest. **d.** Pooled Q eluate fractions. Arrows indicate closely migrating major bands at 150 kDa; asterisks indicate minor bands at 100 and 75 kDa. **e.** Western blot using commercial polyclonal antibody against spike protein. Each lane was loaded with 0.5, 1 or 1.5 µg inhouse-purified WT or Beta S1S2 protein **f.** ACE2-binding assay. Representative curves are shown.

### Physical characterization of ACM-S1S2 + ACM-CpG vaccine

S1S2(WT), S1S2(Beta) and CpG 7909 were separately encapsulated in ACM polymersomes. To quantify the amount of encapsulated protein, polymersomes were lysed using 2.5-5% Triton X-100 and the lysate analyzed using SDS-PAGE against a series of purified protein standards (Fig. 2a). Encapsulated protein content as determined by densitometric measurements were 249.3 µg/ml for S1S2(WT) and 184.8 µg/ml for S1S2(Beta). ACM-CpG was similarly lysed and measured by RP-HPLC. Encapsulated CpG was determined to be 1047.6 µg/ml. An additional preparation of ACM-S1S2(Beta) + ACM-CpG vaccine with enhanced encapsulation (389.1 µg/ml and 2134.8 µg/ml, respectively) was generated to address the volume constraint of IN administration. DLS measurements of ACM-S1S2(WT), ACM-S1S2(Beta), ACM-CpG and final vaccine formulations containing S1S2(WT) or S1S2(Beta) (Fig. 2b) revealed unimodal size distributions with average diameters of 169.4 nm, 174.0 nm, 135.8 nm, 145.0 nm and 146.9 nm, respectively, and polydispersity indices (PDI) < 0.17 consistently.

**Figure 2.**
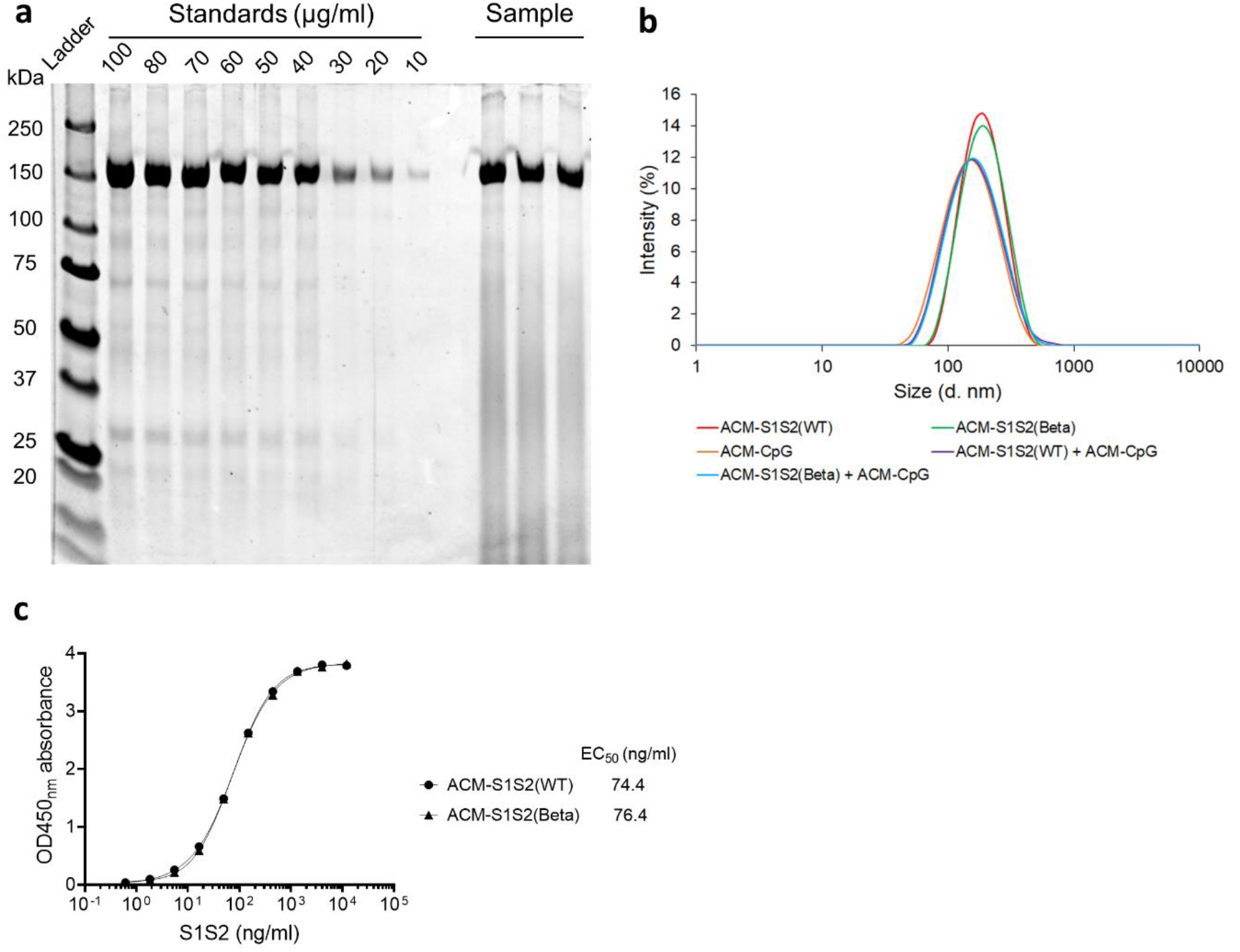
Encapsulation of S1S2 protein within ACM polymersomes. **a.** Quantification of ACM-encapsulated S1S2 protein. Polymersomes were lysed with 2.5% Triton X-100 to release encapsulated material. Pre-diluted ACM lysate was analyzed alongside purified S1S2 standards by SDS-PAGE and protein bands visualized using SYPRO Ruby stain. Concentration of encapsulated S1S2 was determined by densitometry from triplicate measurements. A representative gel image is shown. **b.** Dynamic light scattering (DLS) measurements demonstrating unimodal size distribution of hydrodynamic diameter of ACM-S1S2(WT), ACM-S1S2(Beta), ACM-CpG, and ACM-S1S2 + ACM-CpG formulations. **c.** ACE2-binding assay for ACM-S1S2. Representative curves are shown.

To ensure that protein structure and function were not adversely impacted by encapsulation, ACM-S1S2 was lysed using Triton-X 100 to release protein for ACE2-binding assay. Detergent was then removed using polystyrene adsorbent beads to prevent interference with the assay. Unlike the free protein, S1S2 from the ACM lysate could not be immobilized onto the ELISA plate, possibly due to interference from dissociated ACM polymers. To circumvent this problem, an alternative ELISA format was performed involving the immobilization of ACE2 on the plate followed by titration of S1S2. The assay showed ACM-S1S2(WT) and ACM-S1S2(Beta) to remain functional and bound ACE2 with representative EC_50_ values of 74.4 ng/ml and 76.4 ng/ml, respectively (Fig. 2c).

### ACM-S1S2(Beta) + ACM-CpG induced strong neutralizing titres in golden Syrian hamsters

Two doses of vaccine, containing ACM-Beta or ACM-WT S1S2 protein with ACM-CpG 7909, were administered 21 days apart to hamsters (Fig. 3a). A separate dose-response study in C57BL/6 mice was earlier conducted to identify optimal dose combinations of ACM-S1S2 and ACM-CpG (data not shown). We selected 20 µg S1S2 and 100 µg CpG 7909 for the hamster trial, which also represented the highest intended human dose for our subsequent clinical study. All vaccine formulations were IM administered. In addition, one group of hamsters received ACM-S1S2(Beta) + ACM-CpG via the IN route. Blood and nasal washes were collected on Day 20 (after first dose) and Day 34 (after second dose) to assess antibody response. Spike-specific IgG was clearly detected in serum after a single dose of any vaccine formulation and further increased after the second dose (Fig. 3b, c). Antibodies strongly cross-reacted between WT and Beta spike, which indicated the high degree of identity (99%) between the two proteins. Compared to the free WT spike formulation (i.e.: fS1S2 + fCpG), the ACM-S1S2(WT) + ACM-CpG formulation induced significantly higher IgG after one or two doses, indicating enhancement of immunogenicity through ACM encapsulation. Next, we assessed the immunogenicity of two other formulations: i) an ACM vaccine based on Beta spike protein [i.e.: ACM-S1S2(Beta) + ACM-CpG]; and ii) a bivalent vaccine containing encapsulated WT and Beta spike in a 1:1 ratio [i.e.: ACM-S1S2(WT, Beta) + ACM-CpG]. IM immunization with either vaccine generated comparable IgG responses with geometric mean titers (GMTs) in the range of 10^4^-10^5^ after the second dose, though formulations containing Beta spike did induce slightly higher WT-specific IgG (Fig. 3b). Notably, the ACM-S1S2(Beta) + ACM-CpG formulation was equally immunogenic when administered IM or IN, suggesting that mucosal administration could be a viable vaccination strategy for ACM vaccine formulations.

**Figure 3.**
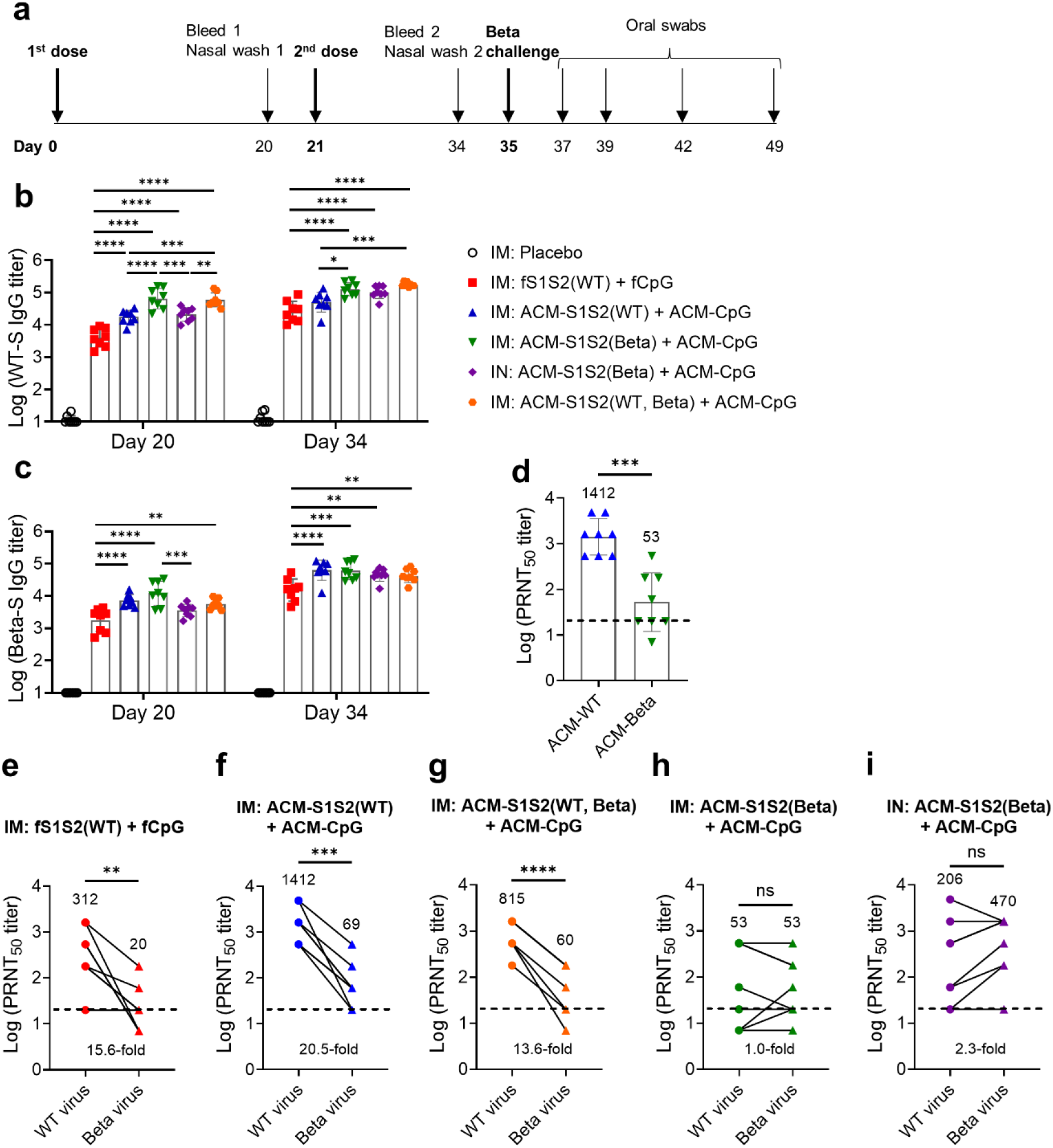
Serum antibody response to ACM-Covid-19 vaccines. **a.** Immunization, sample collection and viral challenge schedule. Golden Syrian hamsters (n = 8) were IM administered one of the following: i) PBS; ii) fS1S2(WT) + fCpG; iii) ACM-S1S2(WT) + ACM-CpG; iv) ACM-S1S2(Beta) + ACM-CpG; v) ACM-S1S2(WT, Beta) + ACM-CpG. In addition, one group of animals received ACM-S1S2(Beta) + ACM-CpG via the IN route. **b, c.** WT and Beta spike-specific IgG titers, respectively. Two-way repeated measures ANOVA (Greenhouse-Geisser correction) with Tukey’s multiple comparison is performed. Only significant differences between vaccinated groups are shown. *: *P* ≤ 0.05; **: *P* ≤ 0.01; ***: *P* ≤ 0.001; ****: *P* ≤ 0.0001. **d.** Homologous PRNT_50_ titers induced by ACM-S1S2(WT) and ACM-S1S2(Beta) vaccines. Day 34 sera were assessed. **e-h.** Neutralizing potency towards WT virus or Beta variant after IM immunization with indicated formulations. **i.** Neutralizing potency towards WT virus or Beta variant after IN immunization with ACM-S1S2(Beta) + ACM-CpG. Geometric mean titers (GMTs) are indicated at the top of each graph; fold-change in GMTs is indicated at the bottom. Two-tailed paired T test is performed.

Serum neutralizing activity was assessed by the plaque reduction neutralization test (PRNT) against SARS-CoV-2 WT virus or Beta variant. First, we compared homologous neutralizing titers to ascertain the relative immunogenicity of each spike protein. ACM-S1S2(WT) + ACM-CpG induced potent neutralizing titers after two doses with a GMT of 1,412 whereas ACM-S1S2(Beta) + ACM-CpG generated modest titers with a GMT of 53 (Fig. 3d), despite similar total IgG titers (Fig 3b, c). With regards to cross-neutralizing ability, antibodies elicited by IM administration of fS1S2(WT) + fCpG, or ACM-S1S2(WT) + ACM-CpG strongly neutralized WT but not Beta virus (Fig. 3e, f). GMT against the Beta variant declined 15-20 folds though majority of hamsters remained seropositive (PRNT_50_ titer ≥ 20). In terms of magnitude, ACM-S1S2(WT) + ACM-CpG vaccine elicited approximately three-fold higher neutralizing titer than the fS1S2(WT) +fCpG formulation, consistent with enhancement of immunogenicity by ACM encapsulation. The bivalent vaccine consisting of ACM-WT and ACM-Beta spike proteins in equal proportion was formulated under the hypothesis that it may induce balanced, cross-neutralizing responses. Unfortunately, a similar sharp drop in neutralizing potency against Beta variant was observed (Fig. 3g). It was previously reported that infection by the Beta variant induced antibodies that strongly cross-neutralized ancestral virus ^30, 31^. Accordingly, hamsters IM vaccinated with the ACM-S1S2(Beta) vaccine generated antibodies that neutralized WT and Beta viruses equally, though the response was modest with a GMT of 53 (Fig. 3h). Remarkably, the same formulation administered via the IN route evoked substantially higher neutralizing titers (GMT 206-470) that remained similarly potent towards WT and Beta viruses (Fig. 3i).

### IN vaccination with ACM-S1S2(Beta) + ACM-CpG strongly protected hamsters from infection and disease by Beta variant

The protective efficacy of each ACM Covid-19 vaccine was assessed using a non-lethal hamster model of Beta infection. Non-vaccinated animals became severely symptomatic and lost up to 13% of body weight on Day 7 post challenge before recovering to their average baseline body weight on Day 14 (Fig. 4a). Animals IM immunized with fS1S2(WT) + fCpG became mildly symptomatic after viral challenge and lost up to 3% of body weight on Day 5 before recovering, suggesting that a serum PRNT50 titer of 20 (Fig. 3e) was associated with partial protection. In contrast, hamsters IM immunized with ACM-S1S2(WT) or ACM-S1S2(Beta) vaccine exhibited progressive weight gain over the 14-day study period (Fig. 4a), suggesting that a serum PRNT_50_ titer >50 (Fig. 3f, h) was associated with full protection against disease. For reasons unclear, the weight gain profile of hamsters IM immunized with the bivalent vaccine [ACM-S1S2(WT, Beta) + ACM-CpG] was slightly attenuated (Fig. 4a) despite an average serum PRNT_50_ titer of 60 (Fig. 3g). Finally, IN administration of ACM-S1S2(Beta) vaccine was also fully protective against weight loss (Fig. 4a), consistent with its robust neutralizing activity towards the Beta variant (Fig. 3i).

**Figure 4.**
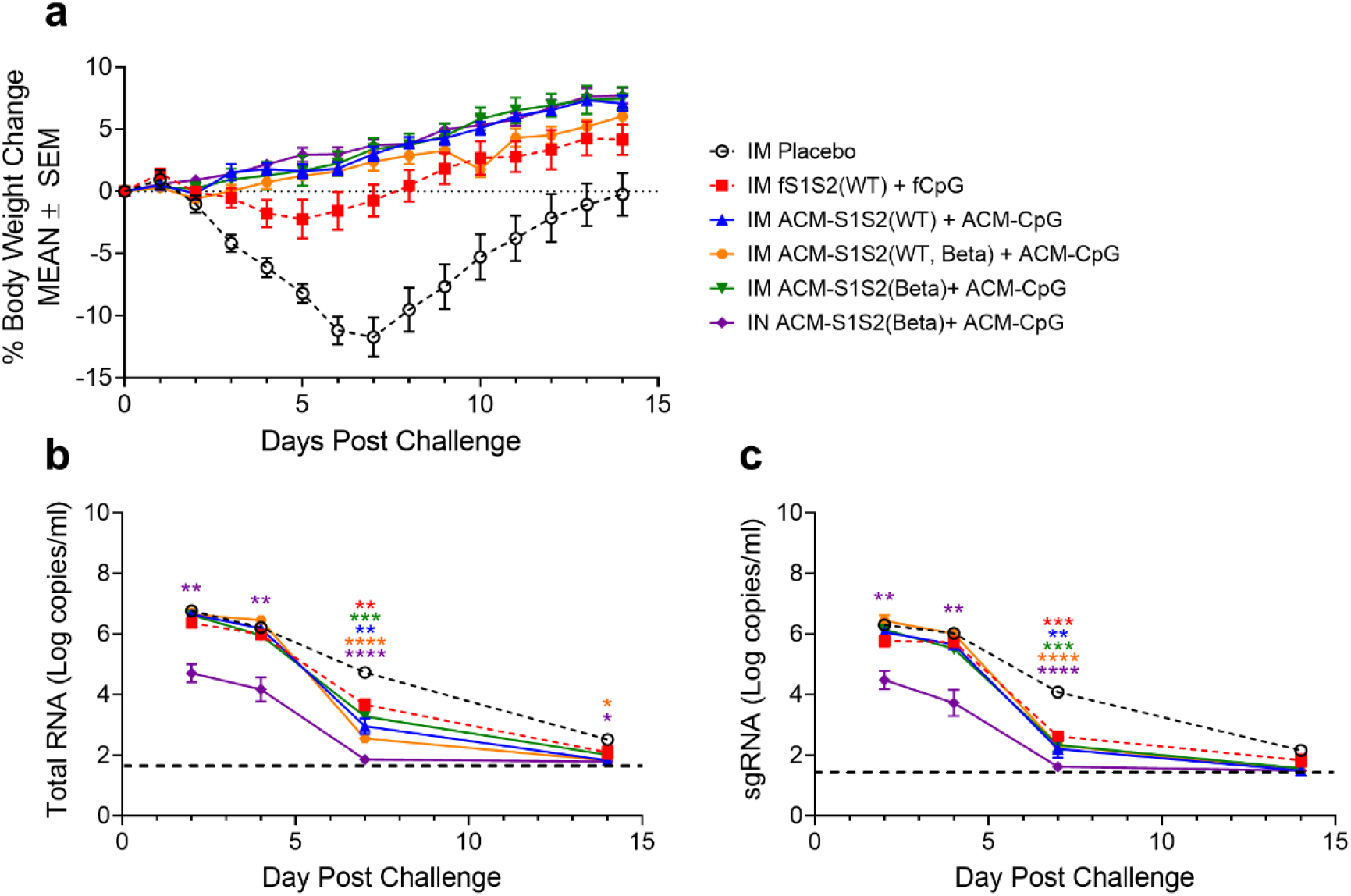
IN challenge with Beta variant. **a.** Animals were observed 14 days for changes in body weight relative to initial (horizontal dashed line). Vaccine formulations and routes of administration are indicated on the right. **b, c.** Viral RNA loads in oral swabs as determined by qPCR. Copy numbers of total and subgenomic RNA (sgRNA) are shown, respectively. Two-way repeated measures ANOVA (Greenhouse-Geisser correction) with Tukey’s multiple comparison is performed. Only significant differences with respect to non-vaccinated controls are shown (color coded). Horizontal dashed lines represent lower limits of detection (62 and 31 RNA copies/ml, respectively).

To assess viral load after infection, oral swabs were collected on Days 2, 4, 7 and 14 (Fig. 3a) and subjected to viral RNA qPCR. Non-vaccinated controls exhibited a peak viral RNA load of 5.8 x 10^6^ copies/ml (geometric mean) on Day 2, which gradually declined to 1.6 x 10^6^ copies/ml on Day 4 and finally 3.3 x 10^2^ copies/ml on Day 14 (Fig. 4b). Hamsters IM immunized with the monovalent ACM-WT spike vaccine, monovalent ACM-Beta spike vaccine, or the bivalent ACM-(WT, Beta) spike vaccine showed similar peak viral loads as the non-vaccinated controls from Days 2 to 4; significant reduction in viral load was seen only on Day 7. In contrast, hamsters IN immunized with ACM-Beta spike vaccine strongly and rapidly suppressed their viral loads. Compared to non-vaccinated controls, the average viral RNA load of IN-vaccinated hamsters was ∼100-folds lower on Days 2 and 4 (5.1 x 10^4^ and 1.5 x 10^4^ RNA copies/ml, respectively) and dropped to the lower limit of detection on Day 7.

To gain further insight into the protective efficacy of ACM vaccines, we performed subgenomic RNA (sgRNA) analysis, which was indicative of viral replication ^41^. In non-vaccinated controls, kinetics of viral sgRNA was similar to total RNA, with peak viral load of 2.0 x 10^6^ copies/ml on Day 2 followed by a decline to 1.1 x 10^6^ copies/ml on Day 4 and finally 1.4 x 10^2^ copies/ml on Day 14 (Fig. 4c). Hamsters IM immunized with any ACM vaccine showed similar sgRNA levels on Days 2 and 4 as the non-vaccinated controls, before a sharp decline on Day 7 and eventually reaching the lower limit of detection on Day 14. In contrast, hamsters IN vaccinated with ACM-S1S2(Beta) + ACM-CpG displayed a 60-200-fold drop in sgRNA on Days 2 to 4 (3.0 x 10^4^ and 5.3 x 10^3^ copies/ml, respectively) and reached the lower limit of detection on Day 7. Taken together, our results indicated that IM vaccination protected against disease but not infection, whereas IN vaccination protected against disease and suppressed viral replication in the upper airway rapidly and potently.

### Antibodies generated by ACM-S1S2(Beta) + ACM-CpG vaccine could neutralize Omicron variant

Since the emergence of Omicron in southern Africa in November 2021, the highly infectious variant has spread rapidly worldwide, even replacing Delta as the dominant strain in certain regions ^23^. It is characterized by more than 30 mutations in the spike protein, located mainly in the NTD and RBD, which are believed to enhance viral fitness and antibody evasion ^19^. Indeed, multiple groups have shown extensive neutralization escape from previously vaccinated or infected subjects ^15, 19, 29, 42–44^. In view of this, we examined nasal washes for antibody response towards Omicron, as an alternative to Day 34 sera which were depleted from earlier tests (i.e., live virus PRNT). For virus neutralizing activity, we utilized an FDA-approved, clinically validated surrogate virus neutralization kit (cPass™) ^45^. Neutralizing antibodies predominantly recognized viral RBD ^3, 4^ and the detection of such antibodies that blocked the interaction between RBD and human ACE2 receptor formed the basis of cPass™. As was previously described by another group ^46^ and from our experience, the highly dilute nature of the nasal wash necessitated the pooling of samples within each group, followed by 40-fold concentration using Vivaspin^®^ columns. Neutralizing activity towards WT, Delta or Omicron virus was not detected in nasal washes from hamsters IM vaccinated with any given formulation (Fig. 5a-d). In contrast, substantial activity towards WT and Delta viruses (65.6% and 61.7% inhibition, respectively) was observed in hamsters IN vaccinated with ACM-S1S2(Beta) + ACM-CpG (Fig. 5e). Neutralizing activity towards Omicron was also detected albeit at a lower level (33.9% inhibition). To confirm these findings, we analyzed sera of C57BL/6 mice IM vaccinated with ACM-S1S2(Beta) + ACM-CpG in a separate dose-response study (selected data described here). Mice that received an optimal dose combination of ACM-S1S2(Beta) + ACM-CpG exhibited a balanced neutralizing profile which retained moderate to high level of activity towards Alpha, Gamma and Delta variants (Fig. 5f). Against Omicron, 2/5 mice had lower activity (30-40% inhibition) whereas the remaining mice displayed high levels of neutralization (70-90% inhibition). Our results therefore suggested that primary vaccination with an optimized dose combination of ACM-S1S2(Beta) + ACM-CpG was sufficient to generate broadly neutralizing antibodies that retained moderate to strong activity towards Omicron. Moreover, immunogenicity was substantially enhanced by IN administration, which also induced neutralizing antibodies in the upper respiratory tract that may suppress an Omicron and, perhaps, future VOC infection.

**Figure 5.**
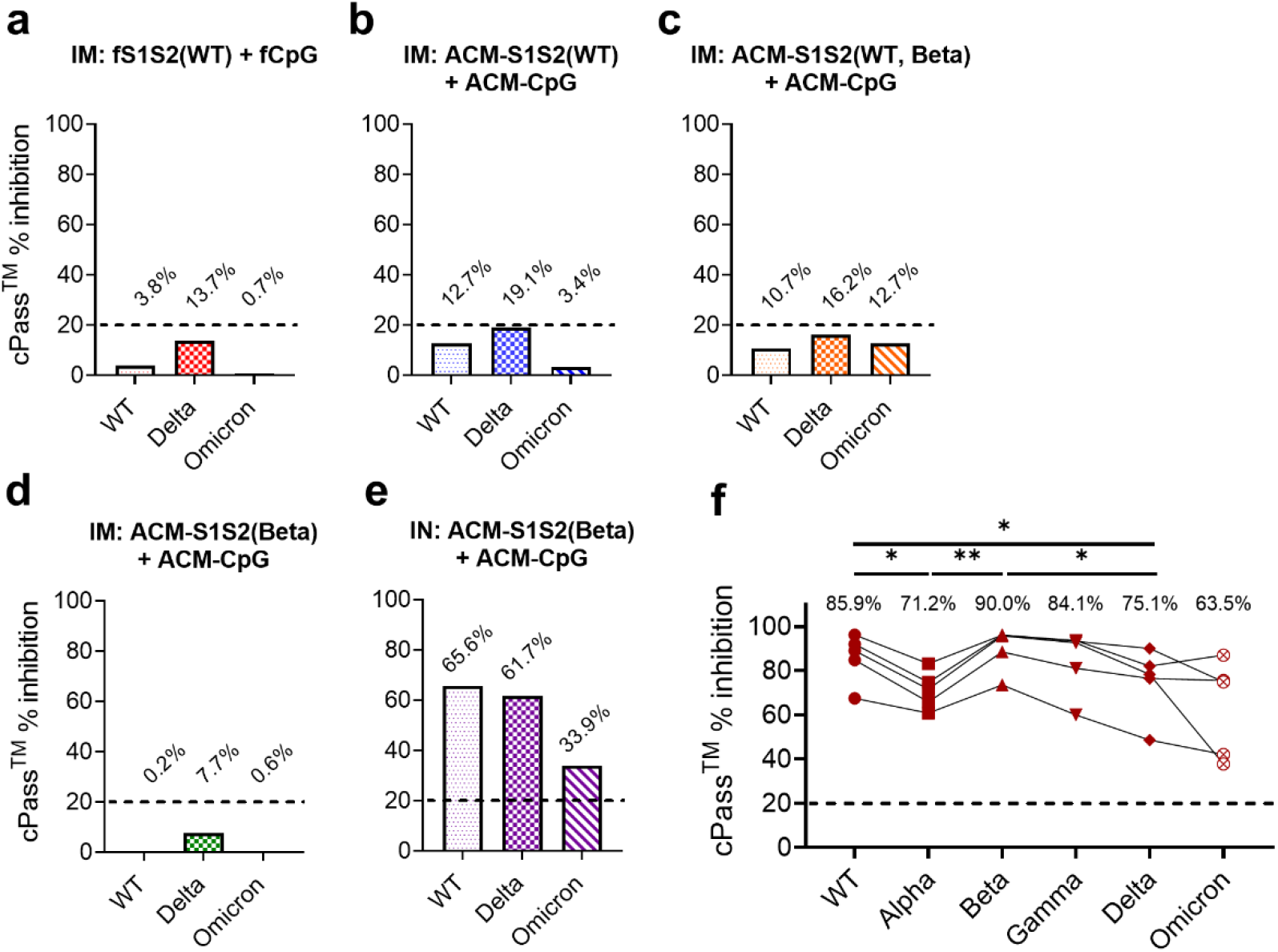
ACM-S1S2(Beta) + ACM-CpG generated Omicron-neutralizing antibody response in hamsters and mice. **a-e.** Virus-neutralizing activity of Day 34 hamster nasal washes. Hamsters were vaccinated with indicated formulations via IM or IN route. To overcome their highly dilute nature, nasal washes from each group were pooled and concentrated 40-folds. Neutralizing activity towards WT, Delta and Omicron viruses were measured using cPass™ surrogate virus neutralization kit (% inhibition indicated above bar graphs). Cutoff of 20% (horizontal dashed line) was recommended by manufacturer to distinguish seropositive samples. **f.** Serum neutralizing activity of C57BL/6 mice (n = 5) IM vaccinated with ACM-S1S2(Beta) + ACM-CpG on Days 0 and 21, from a separate dose-response study. Mouse CpG 1826, instead of human CpG 7909, was used. Sera were examined two weeks after vaccination. Average inhibition of each variant is indicated in graph. One-way repeated measures ANOVA (Greenhouse-Geisser correction) with Tukey’s multiple comparisons is performed – only significant differences are shown. *: *P* ≤ 0.05; **: *P* ≤ 0.01.

## Discussion

We previously reported that ACM polymersomes were efficient vaccine delivery vehicles taken up by DCs and compatible with spike protein and CpG adjuvant to enhance their respective immune responses. Importantly, our Covid-19 vaccine formulation, ACM-S1S2(WT) + ACM-CpG, was highly immunogenic in mice, triggering vigorous antibody and T cell responses ^34^. In the present study, we investigated the ability of ACM vaccines to protect hamsters from a non-lethal SARS-CoV-2 challenge. We chose to focus on the Beta variant because, prior to the emergence of Omicron, it consistently exhibited the strongest potential to evade neutralizing antibodies by existing vaccines ^13, 47, 48^. Notably, a South African clinical trial of the AZD1222 adenovirus vector vaccine established a lack of protection against the Beta variant concomitant with reduced or abrogated antibody neutralization ^49^.

We discovered that the ACM-WT spike vaccine induced markedly higher homologous neutralizing titer than ACM-Beta spike vaccine. A somewhat similar finding was recently reported in which BALB/c immunized with mRNA-1273 (encoding WT spike) generated a pseudovirus neutralizing titer of 16,749 whereas mice immunized with mRNA-1273.351 (encoding Beta spike) had lower titer of 10,948 ^50^. Based on the total IgG response, it was clear that the Beta spike protein was not less immunogenic than WT spike. Since majority of neutralizing antibodies was known to target the RBD and that RBD-reactive antibodies constituted a minor subpopulation of the total spike-specific repertoire ^16^, our results suggested that the immunodominance of Beta RBD may be substantially reduced relative to non-neutralizing or poorly neutralizing epitopes situated elsewhere on the Beta spike. At the same time, we also acknowledged the possibility of a hamster-specific phenomenon arising from species-specific germline antibody genes, since humans infected with the Beta variant could generate similar or stronger neutralizing titers compared to patients infected with WT-like virus^30, 31^.

We established that IM immunization with ACM-WT spike vaccine generated circulating antibodies that potently neutralized WT but not Beta virus, whereas antibodies induced by ACM-Beta spike vaccine neutralized both viruses with similar efficacy. Our hamster findings strongly recapitulated the neutralizing profile of human serum after infection with ancestral or Beta virus ^30, 31^. Changing immunodominance hierarchies may account for such differences. The Beta variant carried ten mutations to the spike protein relative to WT, among which K417N, E484K and N501Y were of particular concern. A recent investigation of the neutralization characteristics of human convalescent plasmas showed that the class 2 neutralizing epitope centered on position 484 was more immunodominant for ancestral SARS-CoV-2, whereas the class 3 epitope spanning sites 443 to 452 was immunodominant for the Beta variant ^51^. This shift in focus towards class 3 epitope may explain the relative insensitivity of Beta immune serum for the mutation at position 484. We also formulated a bivalent ACM vaccine composed of WT and Beta spike in a 1:1 ratio, under the hypothesis that it may elicit cross-neutralizing antibodies. Unfortunately, neutralizing potency towards the Beta variant was similarly reduced, suggesting that WT RBD was more immunogenic and that Beta RBD likely contributed little to the repertoire of neutralizing antibodies. To generate a balanced response with our multivalent formulation, further studies were required to identify the optimal ratio of WT and variant spike.

Although the Beta-specific neutralizing titer induced by IM administration of any ACM vaccine formulation was modest (GMT 53-69), hamsters were fully protected from weight loss after Beta challenge. On the other hand, the fS1S2(WT) + fCpG formulation generated a lower GMT of 20 and hamsters became mildly symptomatic after challenge. These results suggested that a PRNT_50_ titer of 20 may be a protective threshold against severe disease in the context of Beta infection, whereas a PRNT_50_ titer ≥50 may be associated with full protection against disease. Despite the ability of IM vaccination to protect against severe disease, hamsters could not clear the infection efficiently. Peak viral RNA loads from oral swabs were comparable to non-vaccinated controls on Days 2-4 after challenge, and only showed a significant decrease on Day 7. These findings were remarkably similar to the real-world situation, in which IM vaccinated individuals were strongly protected from severe disease and death but substantially less protected against infection ^52^. Studies of Delta variant infection found similar peak viral RNA loads in the upper respiratory tract between vaccinated and non-vaccinated subjects, though vaccination was still associated with a faster decline in viral load ^53, 54^. Poor efficacy against infection was likely due to the inability of parenteral immunization to elicit substantial mucosal immunity, particularly secretory IgA, despite potent systemic immune responses. Mucosal IgA was thought to protect mainly the upper respiratory tract ^55^ and had been shown to neutralize respiratory viruses in humans and animal models, thus serving as a potential correlate of protection ^56, 57^. Conversely, the lack of IgA would result in failure to neutralize SARS-CoV-2 infection of the upper respiratory tract, though systemic immunity remained effective against lower respiratory tract infection and severe disease ^58–60^. With the aim of evoking systemic and mucosal immune responses, we investigated IN administration of ACM-Beta spike vaccine and observed enhanced serum neutralizing titer, potent reduction in viral RNA load of the upper respiratory tract soon after challenge, and virus neutralizing activity in nasal washes. We were unable to measure spike-specific IgA titres, since secondary antibodies targeting hamster IgA were not commercially available. With regards to IM vaccination, although a systemic neutralizing antibody response was generated, mucosal neutralizing activity was not detected and viral replication in the upper respiratory tract was poorly controlled. Cumulatively, our results supported IN vaccination as a relevant strategy to generate a mucosal immune response that effectively inhibited SARS-CoV-2 infection of the upper respiratory tract. Our approach addressed a key weakness of parenteral Covid-19 vaccines and may potentially reduce asymptomatic transmission ^61^.

With more than 30 mutations in its spike protein, Omicron had been shown to extensively escape neutralizing antibodies elicited by past infection or immunization. Primary vaccination with mRNA (BNT162b; mRNA-1273), adenovirus vector (ChAdOx-1 S; Ad26.COV2.S) or subunit protein (NVX-CoV2373) vaccine based on WT spike generated antibodies that poorly neutralized Omicron, leading to complete neutralization escape in 40-100% of vaccinees, depending on the study and the time after immunization ^15, 22, 62^. Nevertheless, neutralizing activity was restored by boosting with mRNA vaccine encoding WT or Omicron spike ^15, 28, 63^. In our preclinical study, we showed that primary vaccination with an optimized dose combination of ACM-S1S2(Beta) and ACM-CpG was sufficient to generate broadly neutralizing antibodies that were efficacious against Delta and Omicron variants. Moreover, neutralizing activity was detected in the upper respiratory tract after IN but not IM vaccination. Enhanced transmissibility of the Omicron variant was thought to be caused, in part, by its robust infection of cells in the upper respiratory tract, compared to the ancestral virus or other variants ^64, 65^. Therefore, the ability to trigger mucosal neutralizing antibodies through IN vaccination could be key to effective control of Omicron infection and may be more relevant than mRNA boosters administered via IM injection.

The main limitation of the present study was that T cell responses had not been investigated, as commercialized antibodies required for assessment of hamster T cell subsets and function were largely unavailable ^66^. Concerns over such constraints of the hamster model had been raised ^67^. Nevertheless, our previous mouse study did establish the presence of functional, memory CD4^+^ and CD8^+^ T cells after primary vaccination ^34^. Unlike neutralizing antibodies, which generally exhibited narrow target specificity as evidenced by moderate to severe reduction in potency against different variants, T cell epitopes were remarkably conserved ^24^. Studies had shown CD4^+^ and CD8^+^ T cells from previous vaccination or infection to retain robust activity against Omicron, despite the variant’s extensive mutations and increased resistance to neutralizing antibodies ^22, 23^. It was believed that cross-reactive T cells may contribute to the control Omicron infection and possibly account for the reduction in disease severity compared to the earlier Delta wave ^68^.

## Acknowledgements

We would like to thank Swagata Kar (PhD) and her team at Bioqual, Inc., U.S.A. for their effort in the hamster experiments and sample analyses, and Joey Poh Soh Yee and the staff of BRC, A*STAR, Singapore for performing mouse immunization and sample collection. We would also like to acknowledge the NHIC Gap Funding Award (NHIC-COV19-2005008) for collaboration with Dr. Francesca Lim, Singapore General Hospital, Singapore.

## Author contributions

The manuscript was written through the contributions of all authors. J.H.L. and M.N. designed the experiments. F.W.I.L. and P.V. provided inputs for the pre-clinical work. D.S., T.W.C., S.L.C., S.V., G.S., T.Y.A., S.W. and J.H.L. performed the experiments. S.K. and the team at Bioqual performed animal experiments and analysis under a paid service agreement. J.H.L., D.S., T.W.C. and M.N. analyzed the data, provided critical intellectual input and wrote the manuscript. All authors have given approval to the final version of this manuscript.

## Conflicting interests

J.H.L., D.S., T.W.C., S.L.C., S.V., G.S., T.Y.A., S.W. and M.N. are employees of ACM Biolabs Pte Ltd, Singapore. P.V is acting Chief Medical Officer of ACM Biosciences AG, Basel, Singapore.

## References

1. Walls, A. C.; Park, Y. J.; Tortorici, M. A.; Wall, A.; McGuire, A. T.; Veesler, D., Structure, Function, and Antigenicity of the SARS-CoV-2 Spike Glycoprotein. Cell 2020, 181 (2), 281–292.e6.

2. Martínez-Flores, D.; Zepeda-Cervantes, J.; Cruz-Reséndiz, A.; Aguirre-Sampieri, S.; Sampieri, A.; Vaca, L., SARS-CoV-2 Vaccines Based on the Spike Glycoprotein and Implications of New Viral Variants. Front Immunol 2021, 12, 701501.

3. Piccoli, L.; Park, Y. J.; Tortorici, M. A.; Czudnochowski, N.; Walls, A. C.; Beltramello, M.; Silacci-Fregni, C.; Pinto, D.; Rosen, L. E.; Bowen, J. E.; Acton, O. J.; Jaconi, S.; Guarino, B.; Minola, A.; Zatta, F.; Sprugasci, N.; Bassi, J.; Peter, A.; De Marco, A.; Nix, J. C.; Mele, F.; Jovic, S.; Rodriguez, B. F.; Gupta, S. V.; Jin, F.; Piumatti, G.; Lo Presti, G.; Pellanda, A. F.; Biggiogero, M.; Tarkowski, M.; Pizzuto, M. S.; Cameroni, E.; Havenar-Daughton, C.; Smithey, M.; Hong, D.; Lepori, V.; Albanese, E.; Ceschi, A.; Bernasconi, E.; Elzi, L.; Ferrari, P.; Garzoni, C.; Riva, A.; Snell, G.; Sallusto, F.; Fink, K.; Virgin, H. W.; Lanzavecchia, A.; Corti, D.; Veesler, D., Mapping Neutralizing and Immunodominant Sites on the SARS-CoV-2 Spike Receptor-Binding Domain by Structure-Guided High-Resolution Serology. Cell 2020, 183 (4), 1024–1042.e21.

4. Greaney, A. J.; Starr, T. N.; Gilchuk, P.; Zost, S. J.; Binshtein, E.; Loes, A. N.; Hilton, S. K.; Huddleston, J.; Eguia, R.; Crawford, K. H. D.; Dingens, A. S.; Nargi, R. S.; Sutton, R. E.; Suryadevara, N.; Rothlauf, P. W.; Liu, Z.; Whelan, S. P. J.; Carnahan, R. H.; Crowe, J. E., Jr.; Bloom, J. D., Complete Mapping of Mutations to the SARS-CoV-2 Spike Receptor-Binding Domain that Escape Antibody Recognition. Cell Host Microbe 2021, 29 (1), 44–57.e9.

5. Noy-Porat, T.; Makdasi, E.; Alcalay, R.; Mechaly, A.; Levy, Y.; Bercovich-Kinori, A.; Zauberman, A.; Tamir, H.; Yahalom-Ronen, Y.; Israeli, M.; Epstein, E.; Achdout, H.; Melamed, S.; Chitlaru, T.; Weiss, S.; Peretz, E.; Rosen, O.; Paran, N.; Yitzhaki, S.; Shapira, S. C.; Israely, T.; Mazor, O.; Rosenfeld, R., A panel of human neutralizing mAbs targeting SARS-CoV-2 spike at multiple epitopes. Nat Commun 2020, 11 (1), 4303.

6. Brouwer, P. J. M.; Caniels, T. G.; van der Straten, K.; Snitselaar, J. L.; Aldon, Y.; Bangaru, S.; Torres, J. L.; Okba, N. M. A.; Claireaux, M.; Kerster, G.; Bentlage, A. E. H.; van Haaren, M. M.; Guerra, D.; Burger, J. A.; Schermer, E. E.; Verheul, K. D.; van der Velde, N.; van der Kooi, A.; van Schooten, J.; van Breemen, M. J.; Bijl, T. P. L.; Sliepen, K.; Aartse, A.; Derking, R.; Bontjer, I.; Kootstra, N. A.; Wiersinga, W. J.; Vidarsson, G.; Haagmans, B. L.; Ward, A. B.; de Bree, G. J.; Sanders, R. W.; van Gils, M. J., Potent neutralizing antibodies from COVID-19 patients define multiple targets of vulnerability. Science 2020, 369 (6504), 643–650.

7. Poh, C. M.; Carissimo, G.; Wang, B.; Amrun, S. N.; Lee, C. Y.; Chee, R. S.; Fong, S. W.; Yeo, N. K.; Lee, W. H.; Torres-Ruesta, A.; Leo, Y. S.; Chen, M. I.; Tan, S. Y.; Chai, L. Y. A.; Kalimuddin, S.; Kheng, S. S. G.; Thien, S. Y.; Young, B. E.; Lye, D. C.; Hanson, B. J.; Wang, C. I.; Renia, L.; Ng, L. F. P., Two linear epitopes on the SARS-CoV-2 spike protein that elicit neutralising antibodies in COVID-19 patients. Nat Commun 2020, 11 (1), 2806.

8. Wang, C.; van Haperen, R.; Gutiérrez-Álvarez, J.; Li, W.; Okba, N. M. A.; Albulescu, I.; Widjaja, I.; van Dieren, B.; Fernandez-Delgado, R.; Sola, I.; Hurdiss, D. L.; Daramola, O.; Grosveld, F.; van Kuppeveld, F. J. M.; Haagmans, B. L.; Enjuanes, L.; Drabek, D.; Bosch, B. J., A conserved immunogenic and vulnerable site on the coronavirus spike protein delineated by cross-reactive monoclonal antibodies. Nat Commun 2021, 12 (1), 1715.

9. Wheatley, A. K.; Pymm, P.; Esterbauer, R.; Dietrich, M. H.; Lee, W. S.; Drew, D.; Kelly, H. G.; Chan, L. J.; Mordant, F. L.; Black, K. A.; Adair, A.; Tan, H. X.; Juno, J. A.; Wragg, K. M.; Amarasena, T.; Lopez, E.; Selva, K. J.; Haycroft, E. R.; Cooney, J. P.; Venugopal, H.; Tan, L. L.; Mt, O. N.; Allison, C. C.; Cromer, D.; Davenport, M. P.; Bowen, R. A.; Chung, A. W.; Pellegrini, M.; Liddament, M. T.; Glukhova, A.; Subbarao, K.; Kent, S. J.; Tham, W. H., Landscape of human antibody recognition of the SARS-CoV-2 receptor binding domain. Cell Rep 2021, 37 (2), 109822.

10. Buchan, S. A.; Tibebu, S.; Daneman, N.; Whelan, M.; Vanniyasingam, T.; Murti, M.; Brown, K. A., Increased household secondary attacks rates with Variant of Concern SARS-CoV-2 index cases. Clin Infect Dis 2021.

11. Dhar, M. S.; Marwal, R.; Vs, R.; Ponnusamy, K.; Jolly, B.; Bhoyar, R. C.; Sardana, V.; Naushin, S.; Rophina, M.; Mellan, T. A.; Mishra, S.; Whittaker, C.; Fatihi, S.; Datta, M.; Singh, P.; Sharma, U.; Ujjainiya, R.; Bhatheja, N.; Divakar, M. K.; Singh, M. K.; Imran, M.; Senthivel, V.; Maurya, R.; Jha, N.; Mehta, P.; A, V.; Sharma, P.; Vr, A.; Chaudhary, U.; Soni, N.; Thukral, L.; Flaxman, S.; Bhatt, S.; Pandey, R.; Dash, D.; Faruq, M.; Lall, H.; Gogia, H.; Madan, P.; Kulkarni, S.; Chauhan, H.; Sengupta, S.; Kabra, S.; Gupta, R. K.; Singh, S. K.; Agrawal, A.; Rakshit, P.; Nandicoori, V.; Tallapaka, K. B.; Sowpati, D. T.; Thangaraj, K.; Bashyam, M. D.; Dalal, A.; Sivasubbu, S.; Scaria, V.; Parida, A.; Raghav, S. K.; Prasad, P.; Sarin, A.; Mayor, S.; Ramakrishnan, U.; Palakodeti, D.; Seshasayee, A. S. N.; Bhat, M.; Shouche, Y.; Pillai, A.; Dikid, T.; Das, S.; Maitra, A.; Chinnaswamy, S.; Biswas, N. K.; Desai, A. S.; Pattabiraman, C.; Manjunatha, M. V.; Mani, R. S.; Arunachal Udupi, G.; Abraham, P.; Atul, P. V.; Cherian, S. S., Genomic characterization and epidemiology of an emerging SARS-CoV-2 variant in Delhi, India. Science 2021, 374 (6570), 995–999.

12. Fisman, D. N.; Tuite, A. R., Evaluation of the relative virulence of novel SARS-CoV-2 variants: a retrospective cohort study in Ontario, Canada. Cmaj 2021, 193 (42), E1619–e1625.

13. Planas, D.; Veyer, D.; Baidaliuk, A.; Staropoli, I.; Guivel-Benhassine, F.; Rajah, M. M.; Planchais, C.; Porrot, F.; Robillard, N.; Puech, J.; Prot, M.; Gallais, F.; Gantner, P.; Velay, A.; Le Guen, J.; Kassis-Chikhani, N.; Edriss, D.; Belec, L.; Seve, A.; Courtellemont, L.; Péré, H.; Hocqueloux, L.; Fafi-Kremer, S.; Prazuck, T.; Mouquet, H.; Bruel, T.; Simon-Lorière, E.; Rey, F. A.; Schwartz, O., Reduced sensitivity of SARS-CoV-2 variant Delta to antibody neutralization. Nature 2021, 596 (7871), 276–280.

14. Liu, C.; Ginn, H. M.; Dejnirattisai, W.; Supasa, P.; Wang, B.; Tuekprakhon, A.; Nutalai, R.; Zhou, D.; Mentzer, A. J.; Zhao, Y.; Duyvesteyn, H. M. E.; López-Camacho, C.; Slon-Campos, J.; Walter, T. S.; Skelly, D.; Johnson, S. A.; Ritter, T. G.; Mason, C.; Costa Clemens, S. A.; Gomes Naveca, F.; Nascimento, V.; Nascimento, F.; Fernandes da Costa, C.; Resende, P. C.; Pauvolid-Correa, A.; Siqueira, M. M.; Dold, C.; Temperton, N.; Dong, T.; Pollard, A. J.; Knight, J. C.; Crook, D.; Lambe, T.; Clutterbuck, E.; Bibi, S.; Flaxman, A.; Bittaye, M.; Belij-Rammerstorfer, S.; Gilbert, S. C.; Malik, T.; Carroll, M. W.; Klenerman, P.; Barnes, E.; Dunachie, S. J.; Baillie, V.; Serafin, N.; Ditse, Z.; Da Silva, K.; Paterson, N. G.; Williams, M. A.; Hall, D. R.; Madhi, S.; Nunes, M. C.; Goulder, P.; Fry, E. E.; Mongkolsapaya, J.; Ren, J.; Stuart, D. I.; Screaton, G. R., Reduced neutralization of SARS-CoV-2 B.1.617 by vaccine and convalescent serum. Cell 2021, 184 (16), 4220- 4236.e13.

15. Garcia-Beltran, W. F.; St Denis, K. J.; Hoelzemer, A.; Lam, E. C.; Nitido, A. D.; Sheehan, M. L.; Berrios, C.; Ofoman, O.; Chang, C. C.; Hauser, B. M.; Feldman, J.; Roederer, A. L.; Gregory, D. J.; Poznansky, M. C.; Schmidt, A. G.; Iafrate, A. J.; Naranbhai, V.; Balazs, A. B., mRNA-based COVID-19 vaccine boosters induce neutralizing immunity against SARS-CoV-2 Omicron variant. Cell 2022.

16. Greaney, A. J.; Loes, A. N.; Crawford, K. H. D.; Starr, T. N.; Malone, K. D.; Chu, H. Y.; Bloom, J. D., Comprehensive mapping of mutations in the SARS-CoV-2 receptor-binding domain that affect recognition by polyclonal human plasma antibodies. Cell Host Microbe 2021, 29 (3), 463–476.e6.

17. Liu, Z.; VanBlargan, L. A.; Bloyet, L. M.; Rothlauf, P. W.; Chen, R. E.; Stumpf, S.; Zhao, H.; Errico, J. M.; Theel, E. S.; Liebeskind, M. J.; Alford, B.; Buchser, W. J.; Ellebedy, A. H.; Fremont, D. H.; Diamond, M. S.; Whelan, S. P. J., Identification of SARS-CoV-2 spike mutations that attenuate monoclonal and serum antibody neutralization. Cell Host Microbe 2021, 29 (3), 477–488.e4.

18. Starr, T. N.; Greaney, A. J.; Hilton, S. K.; Ellis, D.; Crawford, K. H. D.; Dingens, A. S.; Navarro, M. J.; Bowen, J. E.; Tortorici, M. A.; Walls, A. C.; King, N. P.; Veesler, D.; Bloom, J. D., Deep Mutational Scanning of SARS-CoV-2 Receptor Binding Domain Reveals Constraints on Folding and ACE2 Binding. Cell 2020, 182 (5), 1295–1310.e20.

19. Planas, D.; Saunders, N.; Maes, P.; Guivel-Benhassine, F.; Planchais, C.; Buchrieser, J.; Bolland, W. H.; Porrot, F.; Staropoli, I.; Lemoine, F.; Péré, H.; Veyer, D.; Puech, J.; Rodary, J.; Baele, G.; Dellicour, S.; Raymenants, J.; Gorissen, S.; Geenen, C.; Vanmechelen, B.; Wawina-Bokalanga, T.; Martí-Carreras, J.; Cuypers, L.; Sève, A.; Hocqueloux, L.; Prazuck, T.; Rey, F.; Simon-Loriere, E.; Bruel, T.; Mouquet, H.; André, E.; Schwartz, O., Considerable escape of SARS-CoV-2 Omicron to antibody neutralization. Nature 2021.

20. Lopez Bernal, J.; Andrews, N.; Gower, C.; Gallagher, E.; Simmons, R.; Thelwall, S.; Stowe, J.; Tessier, E.; Groves, N.; Dabrera, G.; Myers, R.; Campbell, C. N. J.; Amirthalingam, G.; Edmunds, M.; Zambon, M.; Brown, K. E.; Hopkins, S.; Chand, M.; Ramsay, M., Effectiveness of Covid-19 Vaccines against the B.1.617.2 (Delta) Variant. N Engl J Med 2021, 385 (7), 585-594.

21. Cohn, B. A.; Cirillo, P. M.; Murphy, C. C.; Krigbaum, N. Y.; Wallace, A. W., SARS-CoV-2 vaccine protection and deaths among US veterans during 2021. Science 2021, eabm0620.

22. GeurtsvanKessel, C. H.; Geers, D.; Schmitz, K. S.; Mykytyn, A. Z.; Lamers, M. M.; Bogers, S.; Scherbeijn, S.; Gommers, L.; Sablerolles, R. S. G.; Nieuwkoop, N. N.; Rijsbergen, L. C.; van Dijk, L. L. A.; de Wilde, J.; Alblas, K.; Breugem, T. I.; Rijnders, B. J. A.; de Jager, H.; Weiskopf, D.; van der Kuy, P. H. M.; Sette, A.; Koopmans, M. P. G.; Grifoni, A.; Haagmans, B. L.; de Vries, R. D., Divergent SARS CoV-2 Omicron-reactive T-and B cell responses in COVID-19 vaccine recipients. Sci Immunol 2022, eabo2202.

23. Keeton, R.; Tincho, M. B.; Ngomti, A.; Baguma, R.; Benede, N.; Suzuki, A.; Khan, K.; Cele, S.; Bernstein, M.; Karim, F.; Madzorera, S. V.; Moyo-Gwete, T.; Mennen, M.; Skelem, S.; Adriaanse, M.; Mutithu, D.; Aremu, O.; Stek, C.; Bruyn, E. d.; Van Der Mescht, M. A.; de Beer, Z.; de Villiers, T. R.; Bodenstein, A.; van den Berg, G.; Mendes, A.; Strydom, A.; Venter, M.; Grifoni, A.; Weiskopf, D.; Sette, A.; Wilkinson, R. J.; Bekker, L.-G.; Gray, G.; Ueckermann, V.; Rossouw, T.; Boswell, M. T.; Bihman, J.; Moore, P. L.; Sigal, A.; Ntusi, N. A. B.; Burgers, W. A.; Riou, C., SARS-CoV-2 spike T cell responses induced upon vaccination or infection remain robust against Omicron. medRxiv 2021, 2021.12.26.21268380.

24. Tarke, A.; Sidney, J.; Methot, N.; Yu, E. D.; Zhang, Y.; Dan, J. M.; Goodwin, B.; Rubiro, P.; Sutherland, A.; Wang, E.; Frazier, A.; Ramirez, S. I.; Rawlings, S. A.; Smith, D. M.; da Silva Antunes, R.; Peters, B.; Scheuermann, R. H.; Weiskopf, D.; Crotty, S.; Grifoni, A.; Sette, A., Impact of SARS-CoV-2 variants on the total CD4(+) and CD8(+) T cell reactivity in infected or vaccinated individuals. Cell Rep Med 2021, 2 (7), 100355.

25. Levin, E. G.; Lustig, Y.; Cohen, C.; Fluss, R.; Indenbaum, V.; Amit, S.; Doolman, R.; Asraf, K.; Mendelson, E.; Ziv, A.; Rubin, C.; Freedman, L.; Kreiss, Y.; Regev-Yochay, G., Waning Immune Humoral Response to BNT162b2 Covid-19 Vaccine over 6 Months. N Engl J Med 2021, 385 (24), e84.

26. Naaber, P.; Tserel, L.; Kangro, K.; Sepp, E.; Jürjenson, V.; Adamson, A.; Haljasmägi, L.; Rumm, A. P.; Maruste, R.; Kärner, J.; Gerhold, J. M.; Planken, A.; Ustav, M.; Kisand, K.; Peterson, P., Dynamics of antibody response to BNT162b2 vaccine after six months: a longitudinal prospective study. Lancet Reg Health Eur 2021, 10, 100208.

27. Levine-Tiefenbrun, M.; Yelin, I.; Alapi, H.; Katz, R.; Herzel, E.; Kuint, J.; Chodick, G.; Gazit, S.; Patalon, T.; Kishony, R., Viral loads of Delta-variant SARS-CoV-2 breakthrough infections after vaccination and booster with BNT162b2. Nat Med 2021, 27 (12), 2108–2110.

28. Gagne, M.; Moliva, J. I.; Foulds, K. E.; Andrew, S. F.; Flynn, B. J.; Werner, A. P.; Wagner, D. A.; Teng, I.-T.; Lin, B. C.; Moore, C.; Jean-Baptiste, N.; Carroll, R.; Foster, S. L.; Patel, M.; Ellis, M.; Edara, V.-V.; Maldonado, N. V.; Minai, M.; McCormick, L.; Honeycutt, C. C.; Nagata, B. M.; Bock, K. W.; Dulan, C. N. M.; Cordon, J.; Todd, J.-P. M.; McCarthy, E.; Pessaint, L.; Van Ry, A.; Narvaez, B.; Valentin, D.; Cook, A.; Dodson, A.; Steingrebe, K.; Flebbe, D. R.; Nurmukhambetova, S. T.; Godbole, S.; Henry, A. R.; Laboune, F.; Roberts-Torres, J.; Lorang, C. G.; Amin, S.; Trost, J.; Naisan, M.; Basappa, M.; Willis, J.; Wang, L.; Shi, W.; Doria-Rose, N. A.; Olia, A. S.; Liu, C.; Harris, D. R.; Carfi, A.; Mascola, J. R.; Kwong, P. D.; Edwards, D. K.; Andersen, H.; Lewis, M. G.; Corbett, K. S.; Nason, M. C.; McDermott, A. B.; Suthar, M. S.; Moore, I. N.; Roederer, M.; Sullivan, N. J.; Douek, D. C.; Seder, R. A., mRNA-1273 or mRNA-Omicron boost in vaccinated macaques elicits comparable B cell expansion, neutralizing antibodies and protection against Omicron. bioRxiv 2022, 2022.02.03.479037.

29. Dejnirattisai, W.; Huo, J.; Zhou, D.; Zahradník, J.; Supasa, P.; Liu, C.; Duyvesteyn, H. M. E.; Ginn, H. M.; Mentzer, A. J.; Tuekprakhon, A.; Nutalai, R.; Wang, B.; Dijokaite, A.; Khan, S.; Avinoam, O.; Bahar, M.; Skelly, D.; Adele, S.; Johnson, S. A.; Amini, A.; Ritter, T.; Mason, C.; Dold, C.; Pan, D.; Assadi, S.; Bellass, A.; Omo-Dare, N.; Koeckerling, D.; Flaxman, A.; Jenkin, D.; Aley, P. K.; Voysey, M.; Costa Clemens, S. A.; Naveca, F. G.; Nascimento, V.; Nascimento, F.; Fernandes da Costa, C.; Resende, P. C.; Pauvolid-Correa, A.; Siqueira, M. M.; Baillie, V.; Serafin, N.; Ditse, Z.; Silva, K. D.; Madhi, S.; Nunes, M. C.; Malik, T.; Openshaw, P. J.; Baillie, J. K.; Semple, M. G.; Townsend, A. R.; Huang, K. A.; Tan, T. K.; Carroll, M. W.; Klenerman, P.; Barnes, E.; Dunachie, S. J.; Constantinides, B.; Webster, H.; Crook, D.; Pollard, A. J.; Lambe, T.; Paterson, N. G.; Williams, M. A.; Hall, D. R.; Fry, E. E.; Mongkolsapaya, J.; Ren, J.; Schreiber, G.; Stuart, D. I.; Screaton, G. R., Omicron-B.1.1.529 leads to widespread escape from neutralizing antibody responses. bioRxiv 2021.

30. Cele, S.; Gazy, I.; Jackson, L.; Hwa, S. H.; Tegally, H.; Lustig, G.; Giandhari, J.; Pillay, S.; Wilkinson, E.; Naidoo, Y.; Karim, F.; Ganga, Y.; Khan, K.; Bernstein, M.; Balazs, A. B.; Gosnell, B. I.; Hanekom, W.; Moosa, M. S.; Lessells, R. J.; de Oliveira, T.; Sigal, A., Escape of SARS-CoV-2 501Y.V2 from neutralization by convalescent plasma. Nature 2021, 593 (7857), 142-146.

31. Moyo-Gwete, T.; Madzivhandila, M.; Makhado, Z.; Ayres, F.; Mhlanga, D.; Oosthuysen, B.; Lambson, B. E.; Kgagudi, P.; Tegally, H.; Iranzadeh, A.; Doolabh, D.; Tyers, L.; Chinhoyi, L. R.; Mennen, M.; Skelem, S.; Marais, G.; Wibmer, C. K.; Bhiman, J. N.; Ueckermann, V.; Rossouw, T.; Boswell, M.; de Oliveira, T.; Williamson, C.; Burgers, W. A.; Ntusi, N.; Morris, L.; Moore, P. L., Cross-Reactive Neutralizing Antibody Responses Elicited by SARS-CoV-2 501Y.V2 (B.1.351). N Engl J Med 2021, 384 (22), 2161-2163.

32. Waltz, E., COVID vaccine makers brace for a variant worse than Delta. Nature 2021, 598 (7882), 552–553.

33. Rössler, A.; Riepler, L.; Bante, D.; Laer, D. v.; Kimpel, J., SARS-CoV-2 B.1.1.529 variant (Omicron) evades neutralization by sera from vaccinated and convalescent individuals. medRxiv 2021, 2021.12.08.21267491.

34. Lam, J. H.; Khan, A. K.; Cornell, T. A.; Chia, T. W.; Dress, R. J.; Yeow, W. W. W.; Mohd-Ismail, N. K.; Venkataraman, S.; Ng, K. T.; Tan, Y. J.; Anderson, D. E.; Ginhoux, F.; Nallani, M., Polymersomes as Stable Nanocarriers for a Highly Immunogenic and Durable SARS-CoV-2 Spike Protein Subunit Vaccine. ACS Nano 2021, 15 (10), 15754–15770.

35. Mohsen, M. O.; Gomes, A. C.; Cabral-Miranda, G.; Krueger, C. C.; Leoratti, F. M.; Stein, J. V.; Bachmann, M. F., Delivering adjuvants and antigens in separate nanoparticles eliminates the need of physical linkage for effective vaccination. J Control Release 2017, 251, 92–100.

36. Imai, M.; Iwatsuki-Horimoto, K.; Hatta, M.; Loeber, S.; Halfmann, P. J.; Nakajima, N.; Watanabe, T.; Ujie, M.; Takahashi, K.; Ito, M.; Yamada, S.; Fan, S.; Chiba, S.; Kuroda, M.; Guan, L.; Takada, K.; Armbrust, T.; Balogh, A.; Furusawa, Y.; Okuda, M.; Ueki, H.; Yasuhara, A.; Sakai-Tagawa, Y.; Lopes, T. J. S.; Kiso, M.; Yamayoshi, S.; Kinoshita, N.; Ohmagari, N.; Hattori, S. I.; Takeda, M.; Mitsuya, H.; Krammer, F.; Suzuki, T.; Kawaoka, Y., Syrian hamsters as a small animal model for SARS-CoV-2 infection and countermeasure development. Proc Natl Acad Sci U S A 2020, 117 (28), 16587–16595.

37. Sia, S. F.; Yan, L. M.; Chin, A. W. H.; Fung, K.; Choy, K. T.; Wong, A. Y. L.; Kaewpreedee, P.; Perera, R.; Poon, L. L. M.; Nicholls, J. M.; Peiris, M.; Yen, H. L., Pathogenesis and transmission of SARS-CoV-2 in golden hamsters. Nature 2020, 583 (7818), 834–838.

38. Janakiraman, V.; Forrest, W. F.; Seshagiri, S., Estimation of baculovirus titer based on viable cell size. Nat Protoc 2006, 1 (5), 2271–6.

39. Hsieh, C. L.; Goldsmith, J. A.; Schaub, J. M.; DiVenere, A. M.; Kuo, H. C.; Javanmardi, K.; Le, K. C.; Wrapp, D.; Lee, A. G.; Liu, Y.; Chou, C. W.; Byrne, P. O.; Hjorth, C. K.; Johnson, N. V.; Ludes-Meyers, J.; Nguyen, A. W.; Park, J.; Wang, N.; Amengor, D.; Lavinder, J. J.; Ippolito, G. C.; Maynard, J. A.; Finkelstein, I. J.; McLellan, J. S., Structure-based design of prefusion-stabilized SARS-CoV-2 spikes. Science 2020, 369 (6510), 1501–1505.

40. Riley, T. P.; Chou, H. T.; Hu, R.; Bzymek, K. P.; Correia, A. R.; Partin, A. C.; Li, D.; Gong, D.; Wang, Z.; Yu, X.; Manzanillo, P.; Garces, F., Enhancing the Prefusion Conformational Stability of SARS-CoV-2 Spike Protein Through Structure-Guided Design. Front Immunol 2021, 12, 660198.

41. Wölfel, R.; Corman, V. M.; Guggemos, W.; Seilmaier, M.; Zange, S.; Müller, M. A.; Niemeyer, D.; Jones, T. C.; Vollmar, P.; Rothe, C.; Hoelscher, M.; Bleicker, T.; Brünink, S.; Schneider, J.; Ehmann, R.; Zwirglmaier, K.; Drosten, C.; Wendtner, C., Virological assessment of hospitalized patients with COVID-2019. Nature 2020, 581 (7809), 465–469.

42. Cameroni, E.; Bowen, J. E.; Rosen, L. E.; Saliba, C.; Zepeda, S. K.; Culap, K.; Pinto, D.; VanBlargan, L. A.; De Marco, A.; di Iulio, J.; Zatta, F.; Kaiser, H.; Noack, J.; Farhat, N.; Czudnochowski, N.; Havenar-Daughton, C.; Sprouse, K. R.; Dillen, J. R.; Powell, A. E.; Chen, A.; Maher, C.; Yin, L.; Sun, D.; Soriaga, L.; Bassi, J.; Silacci-Fregni, C.; Gustafsson, C.; Franko, N. M.; Logue, J.; Iqbal, N. T.; Mazzitelli, I.; Geffner, J.; Grifantini, R.; Chu, H.; Gori, A.; Riva, A.; Giannini, O.; Ceschi, A.; Ferrari, P.; Cippà, P. E.; Franzetti-Pellanda, A.; Garzoni, C.; Halfmann, P. J.; Kawaoka, Y.; Hebner, C.; Purcell, L. A.; Piccoli, L.; Pizzuto, M. S.; Walls, A. C.; Diamond, M. S.; Telenti, A.; Virgin, H. W.; Lanzavecchia, A.; Snell, G.; Veesler, D.; Corti, D., Broadly neutralizing antibodies overcome SARS-CoV-2 Omicron antigenic shift. Nature 2021.

43. Cao, Y.; Wang, J.; Jian, F.; Xiao, T.; Song, W.; Yisimayi, A.; Huang, W.; Li, Q.; Wang, P.; An, R.; Wang, J.; Wang, Y.; Niu, X.; Yang, S.; Liang, H.; Sun, H.; Li, T.; Yu, Y.; Cui, Q.; Liu, S.; Yang, X.; Du, S.; Zhang, Z.; Hao, X.; Shao, F.; Jin, R.; Wang, X.; Xiao, J.; Wang, Y.; Xie, X. S., Omicron escapes the majority of existing SARS-CoV-2 neutralizing antibodies. Nature 2021.

44. Cele, S.; Jackson, L.; Khoury, D. S.; Khan, K.; Moyo-Gwete, T.; Tegally, H.; San, J. E.; Cromer, D.; Scheepers, C.; Amoako, D. G.; Karim, F.; Bernstein, M.; Lustig, G.; Archary, D.; Smith, M.; Ganga, Y.; Jule, Z.; Reedoy, K.; Hwa, S. H.; Giandhari, J.; Blackburn, J. M.; Gosnell, B. I.; Abdool Karim, S. S.; Hanekom, W.; von Gottberg, A.; Bhiman, J. N.; Lessells, R. J.; Moosa, M. S.; Davenport, M. P.; de Oliveira, T.; Moore, P. L.; Sigal, A., Omicron extensively but incompletely escapes Pfizer BNT162b2 neutralization. Nature 2021.

45. Tan, C. W.; Chia, W. N.; Qin, X.; Liu, P.; Chen, M. I.; Tiu, C.; Hu, Z.; Chen, V. C.; Young, B. E.; Sia, W. R.; Tan, Y. J.; Foo, R.; Yi, Y.; Lye, D. C.; Anderson, D. E.; Wang, L. F., A SARS-CoV-2 surrogate virus neutralization test based on antibody-mediated blockage of ACE2-spike protein-protein interaction. Nat Biotechnol 2020, 38 (9), 1073–1078.

46. Okamoto, S.; Matsuoka, S.; Takenaka, N.; Haredy, A. M.; Tanimoto, T.; Gomi, Y.; Ishikawa, T.; Akagi, T.; Akashi, M.; Okuno, Y.; Mori, Y.; Yamanishi, K., Intranasal immunization with a formalin-inactivated human influenza A virus whole-virion vaccine alone and intranasal immunization with a split-virion vaccine with mucosal adjuvants show similar levels of cross-protection. Clin Vaccine Immunol 2012, 19 (7), 979–90.

47. Wall, E. C.; Wu, M.; Harvey, R.; Kelly, G.; Warchal, S.; Sawyer, C.; Daniels, R.; Hobson, P.; Hatipoglu, E.; Ngai, Y.; Hussain, S.; Nicod, J.; Goldstone, R.; Ambrose, K.; Hindmarsh, S.; Beale, R.; Riddell, A.; Gamblin, S.; Howell, M.; Kassiotis, G.; Libri, V.; Williams, B.; Swanton, C.; Gandhi, S.; Bauer, D. L., Neutralising antibody activity against SARS-CoV-2 VOCs B.1.617.2 and B.1.351 by BNT162b2 vaccination. Lancet 2021, 397 (10292), 2331-2333.

48. Garcia-Beltran, W. F.; Lam, E. C.; St Denis, K.; Nitido, A. D.; Garcia, Z. H.; Hauser, B. M.; Feldman, J.; Pavlovic, M. N.; Gregory, D. J.; Poznansky, M. C.; Sigal, A.; Schmidt, A. G.; Iafrate, A. J.; Naranbhai, V.; Balazs, A. B., Multiple SARS-CoV-2 variants escape neutralization by vaccine-induced humoral immunity. Cell 2021, 184 (9), 2523.

49. Madhi, S. A.; Baillie, V.; Cutland, C. L.; Voysey, M.; Koen, A. L.; Fairlie, L.; Padayachee, S. D.; Dheda, K.; Barnabas, S. L.; Bhorat, Q. E.; Briner, C.; Kwatra, G.; Ahmed, K.; Aley, P.; Bhikha, S.; Bhiman, J. N.; Bhorat, A. E.; du Plessis, J.; Esmail, A.; Groenewald, M.; Horne, E.; Hwa, S. H.; Jose, A.; Lambe, T.; Laubscher, M.; Malahleha, M.; Masenya, M.; Masilela, M.; McKenzie, S.; Molapo, K.; Moultrie, A.; Oelofse, S.; Patel, F.; Pillay, S.; Rhead, S.; Rodel, H.; Rossouw, L.; Taoushanis, C.; Tegally, H.; Thombrayil, A.; van Eck, S.; Wibmer, C. K.; Durham, N. M.; Kelly, E. J.; Villafana, T. L.; Gilbert, S.; Pollard, A. J.; de Oliveira, T.; Moore, P. L.; Sigal, A.; Izu, A., Efficacy of the ChAdOx1 nCoV-19 Covid-19 Vaccine against the B.1.351 Variant. N Engl J Med 2021, 384 (20), 1885-1898.

50. Wu, K.; Choi, A.; Koch, M.; Elbashir, S.; Ma, L.; Lee, D.; Woods, A.; Henry, C.; Palandjian, C.; Hill, A.; Jani, H.; Quinones, J.; Nunna, N.; O’Connell, S.; McDermott, A. B.; Falcone, S.; Narayanan, E.; Colpitts, T.; Bennett, H.; Corbett, K. S.; Seder, R.; Graham, B. S.; Stewart-Jones, G. B. E.; Carfi, A.; Edwards, D. K., Variant SARS-CoV-2 mRNA vaccines confer broad neutralization as primary or booster series in mice. Vaccine 2021, 39 (51), 7394–7400.

51. Greaney, A. J.; Starr, T. N.; Eguia, R. T.; Loes, A. N.; Khan, K.; Karim, F.; Cele, S.; Bowen, J. E.; Logue, J. K.; Corti, D.; Veesler, D.; Chu, H. Y.; Sigal, A.; Bloom, J. D., A SARS-CoV-2 variant elicits an antibody response with a shifted immunodominance hierarchy. bioRxiv 2021.

52. Krause, P. R.; Fleming, T. R.; Peto, R.; Longini, I. M.; Figueroa, J. P.; Sterne, J. A. C.; Cravioto, A.; Rees, H.; Higgins, J. P. T.; Boutron, I.; Pan, H.; Gruber, M. F.; Arora, N.; Kazi, F.; Gaspar, R.; Swaminathan, S.; Ryan, M. J.; Henao-Restrepo, A. M., Considerations in boosting COVID-19 vaccine immune responses. Lancet 2021, 398 (10308), 1377–1380.

53. Singanayagam, A.; Hakki, S.; Dunning, J.; Madon, K. J.; Crone, M. A.; Koycheva, A.; Derqui-Fernandez, N.; Barnett, J. L.; Whitfield, M. G.; Varro, R.; Charlett, A.; Kundu, R.; Fenn, J.; Cutajar, J.; Quinn, V.; Conibear, E.; Barclay, W.; Freemont, P. S.; Taylor, G. P.; Ahmad, S.; Zambon, M.; Ferguson, N. M.; Lalvani, A., Community transmission and viral load kinetics of the SARS-CoV-2 delta (B.1.617.2) variant in vaccinated and unvaccinated individuals in the UK: a prospective, longitudinal, cohort study. Lancet Infect Dis 2021.

54. Chia, P. Y.; Ong, S. W. X.; Chiew, C. J.; Ang, L. W.; Chavatte, J. M.; Mak, T. M.; Cui, L.; Kalimuddin, S.; Chia, W. N.; Tan, C. W.; Chai, L. Y. A.; Tan, S. Y.; Zheng, S.; Lin, R. T. P.; Wang, L.; Leo, Y. S.; Lee, V. J.; Lye, D. C.; Young, B. E., Virological and serological kinetics of SARS-CoV-2 Delta variant vaccine breakthrough infections: a multicentre cohort study. Clin Microbiol Infect 2021.

55. Krammer, F., SARS-CoV-2 vaccines in development. Nature 2020, 586 (7830), 516–527.

56. See, R. H.; Zakhartchouk, A. N.; Petric, M.; Lawrence, D. J.; Mok, C. P. Y.; Hogan, R. J.; Rowe, T.; Zitzow, L. A.; Karunakaran, K. P.; Hitt, M. M.; Graham, F. L.; Prevec, L.; Mahony, J. B.; Sharon, C.; Auperin, T. C.; Rini, J. M.; Tingle, A. J.; Scheifele, D. W.; Skowronski, D. M.; Patrick, D. M.; Voss, T. G.; Babiuk, L. A.; Gauldie, J.; Roper, R. L.; Brunham, R. C.; Finlay, B. B., Comparative evaluation of two severe acute respiratory syndrome (SARS) vaccine candidates in mice challenged with SARS coronavirus. J Gen Virol 2006, 87 (Pt 3), 641–650.

57. Morokutti, A.; Muster, T.; Ferko, B., Intranasal vaccination with a replication-deficient influenza virus induces heterosubtypic neutralising mucosal IgA antibodies in humans. Vaccine 2014, 32 (17), 1897–900.

58. Lapuente, D.; Fuchs, J.; Willar, J.; Vieira Antão, A.; Eberlein, V.; Uhlig, N.; Issmail, L.; Schmidt, A.; Oltmanns, F.; Peter, A. S.; Mueller-Schmucker, S.; Irrgang, P.; Fraedrich, K.; Cara, A.; Hoffmann, M.; Pöhlmann, S.; Ensser, A.; Pertl, C.; Willert, T.; Thirion, C.; Grunwald, T.; Überla, K.; Tenbusch, M., Protective mucosal immunity against SARS-CoV-2 after heterologous systemic prime-mucosal boost immunization. Nat Commun 2021, 12 (1), 6871.

59. Hassan, A. O.; Shrihari, S.; Gorman, M. J.; Ying, B.; Yaun, D.; Raju, S.; Chen, R. E.; Dmitriev, I. P.; Kashentseva, E.; Adams, L. J.; Mann, C.; Davis-Gardner, M. E.; Suthar, M. S.; Shi, P. Y.; Saphire, E. O.; Fremont, D. H.; Curiel, D. T.; Alter, G.; Diamond, M. S., An intranasal vaccine durably protects against SARS-CoV-2 variants in mice. Cell Rep 2021, 36 (4), 109452.

60. van Doremalen, N.; Purushotham, J. N.; Schulz, J. E.; Holbrook, M. G.; Bushmaker, T.; Carmody, A.; Port, J. R.; Yinda, C. K.; Okumura, A.; Saturday, G.; Amanat, F.; Krammer, F.; Hanley, P. W.; Smith, B. J.; Lovaglio, J.; Anzick, S. L.; Barbian, K.; Martens, C.; Gilbert, S. C.; Lambe, T.; Munster, V. J., Intranasal ChAdOx1 nCoV-19/AZD1222 vaccination reduces viral shedding after SARS-CoV-2 D614G challenge in preclinical models. Sci Transl Med 2021, 13 (607).

61. Marc, A.; Kerioui, M.; Blanquart, F.; Bertrand, J.; Mitjà, O.; Corbacho-Monné, M.; Marks, M.; Guedj, J., Quantifying the relationship between SARS-CoV-2 viral load and infectiousness. Elife 2021, 10.

62. Mallory, R.; Formica, N.; Pfeiffer, S.; Wilkinson, B.; Marcheschi, A.; Albert, G.; McFall, H.; Robinson, M.; Plested, J. S.; Zhu, M.; Cloney-Clark, S.; Zhou, B.; Chau, G.; Robertson, A.; Maciejewski, S.; Smith, G.; Patel, N.; Glenn, G. M.; Dubovsky, F.; Group, f. t. N. I. n.-S., Immunogenicity and Safety Following a Homologous Booster Dose of a SARS-CoV-2 recombinant spike protein vaccine (NVX-CoV2373): A Phase 2 Randomized Placebo-Controlled Trial. medRxiv 2021, 2021.12.23.21267374.

63. Xia, H.; Zou, J.; Kurhade, C.; Cai, H.; Yang, Q.; Cutler, M.; Cooper, D.; Muik, A.; Jansen, K. U.; Xie, X.; Swanson, K. A.; Shi, P.-Y., Neutralization of Omicron SARS-CoV-2 by 2 or 3 doses of BNT162b2 vaccine. bioRxiv 2022, 2022.01.21.476344.

64. McMahan, K.; Giffin, V.; Tostanoski, L. H.; Chung, B.; Siamatu, M.; Suthar, M. S.; Halfmann, P.; Kawaoka, Y.; Piedra-Mora, C.; Martinot, A. J.; Kar, S.; Andersen, H.; Lewis, M. G.; Barouch, D. H., Reduced Pathogenicity of the SARS-CoV-2 Omicron Variant in Hamsters. bioRxiv 2022, 2022.01.02.474743.

65. Peacock, T. P.; Brown, J. C.; Zhou, J.; Thakur, N.; Newman, J.; Kugathasan, R.; Sukhova, K.; Kaforou, M.; Bailey, D.; Barclay, W. S., The SARS-CoV-2 variant, Omicron, shows rapid replication in human primary nasal epithelial cultures and efficiently uses the endosomal route of entry. bioRxiv 2022, 2021.12.31.474653.

66. Horiuchi, S.; Oishi, K.; Carrau, L.; Frere, J.; Møller, R.; Panis, M.; tenOever, B. R., Immune memory from SARS-CoV-2 infection in hamsters provides variant-independent protection but still allows virus transmission. Sci Immunol 2021, 6 (66), eabm3131.

67. Rees, J.; Haig, D.; Mack, V.; Davis, W. C., Characterisation of monoclonal antibodies specific for hamster leukocyte differentiation molecules. Vet Immunol Immunopathol 2017, 183, 40–44.

68. Wolter, N.; Jassat, W.; Walaza, S.; Welch, R.; Moultrie, H.; Groome, M.; Amoako, D. G.; Everatt, J.; Bhiman, J. N.; Scheepers, C.; Tebeila, N.; Chiwandire, N.; du Plessis, M.; Govender, N.; Ismail, A.; Glass, A.; Mlisana, K.; Stevens, W.; Treurnicht, F. K.; Makatini, Z.; Hsiao, N.-y.; Parboosing, R.; Wadula, J.; Hussey, H.; Davies, M.-A.; Boulle, A.; von Gottberg, A.; Cohen, C., Early assessment of the clinical severity of the SARS-CoV-2 omicron variant in South Africa: a data linkage study. The Lancet.

